# Characterizing the Effects of Glutathione as an Immunoadjuvant in the Treatment of Tuberculosis

**DOI:** 10.1101/335323

**Authors:** Ruoqiong Cao, Garrett Teskey, Hicret Islamoglu, Rachel Abrahem, Karo Gyurjian, Li Zhong, Vishwanath Venketaraman

## Abstract

Mycobacterium *tuberculosis* (*M*. *tb*) is the etiological agent that is responsible for causing tuberculosis (TB), which continues to affect millions of people worldwide, with an ever-increasing resistance to antibiotics. We tested the synergistic effects of N-acetyl cysteine (NAC, the precursor molecule for the synthesis of glutathione) and individual first-line antibiotics typically given for the treatment of TB such as Isoniazid (INH), Rifampicin (RIF), Ethambutol (EMB) and Pyrazinamide (PZA) to improve the ability of macrophages to control intracellular *M*. *tb* infection. Glutathione (GSH), a pleiotropic antioxidant molecule has been previously shown to display both antimycobacterial and immune-enhancing effects. Our results indicate that there was not only an increase in beneficial immunomodulatory effects, but a greater reduction in the intracellular viability of *M. tb* when macrophages were treated with the combination of antibiotics (INH/RIF/EMB or PZA) and NAC.

## Introduction

*Mycobacterium tuberculosis* (*M*. *tb*), is the causative agent of tuberculosis (TB), and a leading cause of death worldwide [1]. According to the World Health Organization (WHO), TB is currently the ninth leading cause of mortality worldwide and the principal cause of death due to a single infectious agent [2]. The WHO reported that 10.4 million people contracted an active TB infection in 2016 alone [2]. *M*. *tb* infection is acquired via inhalation of respiratory droplets, leading to the seeding of the bacteria within the lungs. More specifically, once *M*. *tb* enters the lower respiratory tract, it is engulfed by alveolar macrophages and becomes an intracellular pathogen [3]. At this point a competent immune system will mount an attack against the bacteria, either killing it off completely, or more characteristically, sequestering the bacteria within a specialized immune structure known as a granuloma, archetypally localized within the lungs [4]. This process of mycobacterial containment within a granuloma is referred to as latent TB and is observed in the majority of TB cases. A granuloma consists of a multitude of immune cells which come together to orcistate this protective immune response, including: macrophages, epithelioid histiocytes, dendritic cells, T cells and natural killer (NK) cells, which isolate the bacteria rendering it latent [5]. The accumulation of these immune cells is mediated by various cytokines, including TNF-α, IL-6, IL-12, IL-2 and IFN-γ[6].

However, in an immunocompromised individual *M. tb* granulomas can undergo liquefaction resulting in an active TB infection [7]. Individuals with active TB are not only contagious but are at a serious risk for developing permanent morbidity due to vast cellular damage. Therefore, the Center for Disease Control and Prevention (CDC) recommends that the preferred treatment regimen for an active TB consists of the combined administration of the antibiotics Isoniazid (INH), Rifampicin (RIF), Pyrazinamide (PZA), and Ethambutol (EMB) for 2 months (the intensive phase), followed by the administration of INH and RIF for 4 months (the continuation phase) [8-10]. These antibiotics are primarily used in combination to prevent the bacteria from developing resistance. In lieu of RIF, rifapentine may be used for intermittent dosing, or rifabutin as it has fewer interactions with anti-HIV medications and opioid substitution therapy [11]. However, these alternative medications are more expensive than RIF and are not universally available in all TB programs [12]. Positive prognostic indicators include patients feeling better within a few weeks of beginning treatment and will typically become non-infectious during the intensive phase [12]. The intensive phase is followed by the continuation phase of treatment which consists of four-months of treatment with INH and RIF (the two most powerful first-line antibiotics) [8-10]. A patient is declared cured upon completing this treatment and achieving two negative TB tests. However, TB treatment has been experiencing a rising risk of failure due to the development of multidrug resistant strains of *M*. *tb*, largely due to noncompliance and inadequate adherence to standard treatment protocols and the necessary cessation of treatment due to the detrimental side effects that can be associated with the use of the aforementioned antibiotics [13]. Side effects, particularly nausea and abdominal pain, are relatively common [14]. Additionally, urine and tears can turn orange, which is harmless but can be disconcerting if patients are not forewarned [15]. More severe side effects, such as joint pain, visual impairment, peripheral neuropathy (nerve damage) and hepatotoxicity leading to liver damage are less common but can be serious when they do occur [16]. In 2012, bedaquiline, used to treat drug resistant-TB (DR-TB), became the first new TB drug from a new class to be approved by the U.S. Food and Drug Administration (FDA) in over 40 years [17]. In 2014, bedaquiline’s U.S. accelerated approval was followed by the European Medicines Agency’s (EMA’s) conditional approval of bedaquiline as well as another new drug, delamanid, for the treatment of specific forms of DR-TB, emphasizing the urgent need to develop new treatment options for both drug sensitive and drug resistant TB [18].

In search of a novel therapeutic agent to augment the treatment of TB, we investigated the synergistic effects of the glutathione (GSH) precursor N-acetyl cysteine (NAC), in conjunction with the individual aforementioned first-line antibiotics in promoting macrophage-mediated killing of *M*. *tb*. NAC has been widely used for several years to enhance the intracellular levels of GSH [19]. GSH, a tripeptide comprised of glutamate, cysteine and glycine, functions to protect the cells and tissues from oxidative damage thereby restoring redox homeostasis within the body [20]. Further studies have demonstrated that the biological antioxidant GSH has both antimycobacterial effects and immune-modulating properties [21-23].

Our laboratory has previously reported that the levels of GSH were significantly decreased in the red blood cells, NK cells, macrophages and T cells derived from the peripheral blood of individuals with HIV infection [24-26]. Furthermore, we have also demonstrated that the levels of GSH were significantly compromised in brain tissue samples derived from the frontal cortex of individuals with HIV infection [21]. The decreased levels of GSH among the HIV infected individuals correlated with the diminished control of *M. tb* infection [21-23]. Additional studies have shown that the levels of GSH were significantly compromised in individuals with type 2 diabetes (T2DM) due to diminished levels of enzymes involved in the GSH synthesis [27]. Findings from our clinical trials indicate that supplementation with liposomal glutathione (L-GSH) restored redox homeostasis, induced cytokine balance and improved immune responses against M. *tb* infection [23]. Additionally, NAC has also been shown to have direct mycobactericidal effects [28].

Therefore, we hypothesized that treatment of THP-1 cells with NAC in conjunction with any one of the first line antibiotics: INH, RIF, EMB or PZA will improve the ability of macrophages to effectively control *M. tb* infection. Our study findings indicate a greater reduction in the intracellular viability of *M*. *tb* when macrophages were treated with the combination of NAC and antibiotics (INH/RIF/EMB or PZA).

## Material and Methods

### THP-1 cell culture

THP-1 cells were maintained RPMI media (Sigma) containing 10% Fetal bovine serum (FBS-Sigma) and 2mM glutamine (Sigma). For the assays, cell suspension was centrifuged at 2,000 rpm for 15 minutes, and the pellet was resuspended in RPMI containing 10% FBS. Cell numbers in the suspension were determined by trypan blue dye exclusion staining. THP-1 cells (2 × 10^5^ cells/well) were distributed in 24-well plates (Corning), treated with PMA (Phorbol 12-myristate 13-acetate) at 10 ng/ml concentration and incubated overnight at 37°C, 5% CO2 to induce the differentiation to macrophages. Following overnight incubation, media in the wells were replaced with fresh media.

### Preparation of bacteria for infection assays

The Erdman strain of *M. tb* (was gifted by Dr. Selvakumar Subbian. Rutgers New Jersey Medical School, Biomedical and Health Sciences) was used for all our infection studies. The Erdman strain of M. *tb* (will henceforth be referred to as *M. tb*) has slightly faster doubling time and is more virulent compared to the standard laboratory strain, H37Rv [29]. *M. tb* was cultured in 7H9 media (Middlebrook) supplemented with albumin dextrose complex (ADC) at 37°C until processing. The *M. tb* was processed for infection once the static culture was at the peak logarithmic phase of growth (OD between 0.5-0.8 at A600), and subsequently washed and resuspended in sterile 1X PBS. Bacterial clumps were dispersed by vortexing five times with 3mm sterile glass beads at two minutes intervals. The bacterial suspension was then filtered using a 5μm syringe filter (Millipore) to remove any remaining bacterial aggregations. The single cell suspension of processed *M. tb* was serially diluted and plated on 7H11 agar to determine the bacterial numbers in the processed stock. Aliquots of processed bacterial stocks were frozen at −80°C. At the time of infection, the processed frozen stocks of *M*. *tb* were thawed and used for the infection. All infection studies and handling of the *M. tb* was done inside a certified biosafety level 3 facility (BSL-3).

### THP-1 macrophage infection and treatment

Differentiated macrophages were infected with processed *M. tb* at a multiplicity of infection or MOI of 0.1:1 (bacteria to macrophage ratio). Infected macrophages were incubated for 1 hour and then successively washed 3 times with warm 1X PBS to remove the unphagocytozed bacteria. Infected macrophages were then either sham-treated or received a onetime treatment with the MIC of the respective antibiotic with or without NAC addition. NAC (10 mM) was administered at 3 equal intervals throughout the trial. The treatment concentrations administered were as follows: INH (0.125 micrograms/ml), INH (0.125 micrograms/ml) + NAC (10 mM (x3)), RIF (0.125 micrograms/ml), RIF (0.125 micrograms/ml) + NAC (10 mM (x3)), EMB (8.0 micrograms/ml), EMB (8.0 micrograms/ml) + NAC (10 mM (x3)), PZA (50 micrograms/ml) and PZA (50 micrograms/ml) + NAC (10 mM (x3)). The Infected cells were maintained at 37°C, 5% CO2 until they were terminated at 1 hour and 12 days post-infection to determine the intracellular survival of M. *tb*.

### Termination of macrophages and CFUs assay

Termination of infected macrophages was performed by collecting and storing the cell-free supernatants and lysing THP-1 cells using 250 ul of ice cold, sterile 1X PBS. Cell lysates collected from the wells were vigorously vortexed and then subjected to freeze/thaw cycles to ensure complete lysis of macrophages. The collected lysates and supernatants were then diluted in sterile 1X PBS and plated on 7H11 medium (Hi Media) enriched with ADC to evaluate M. *tb* survival inside the macrophages by counting the bacterial colonies.

### Quantification of GSH levels in the cellular lysates

The quantity of total glutathione present was measured by the colorimetric method using an assay kit from Arbor Assay (K006-H1). The macrophages lysates were first thoroughly mixed with an equal volume of cold 5% sulfosalicylic acid (SSA), and then incubated for 10 minutes at 4°C, followed by centrifugation at 14000 rpm for 10 minutes. The GSH was measured following the manufacturer’s instructions. All measurements were normalized by total protein levels and the results were reported in moles of GSH per gram of protein.

### Cytokines measurements of culture medium

The effects of *M. tb* infection, antibiotic and NAC treatments in altering the production of TNF-α and IL-10 was determined by quantifying the levels of these cytokines in the macrophage supernatants collected at 12 days post-infection by enzyme-linked immunosorbent assay (ELISA) using assay kits from Affymetrix as per the manufacturer’s protocol..

### Statistical Analysis

Statistical data analysis was performed using GraphPad Prism Software version 7. Levels of cytokines, GSH, MDA and CFUs were compared between untreated control, antibiotic-treatment and treatment with antibiotics in conjunction with NAC using the unpaired t-test with Welch correction. Reported values are in means. P<0.05 was considered significant (*p<0.05, **p<0.005).

## Results

### Levels of total GSH in uninfected and *M. tb*-infected THP-1 cells

GSH levels were measured in the lysates of THP-1 cells using an assay kit from Arbor Assays. The levels of total GSH were shown to be significantly diminished in *M*. *tb* infected macrophages compared to the uninfected control group (Fig. 1A). These results indicate that infection with Erdman strain of *M. tb* can cause a significant two-fold decrease in the intracellular levels of GSH in macrophages.

**Fig. 1A:**
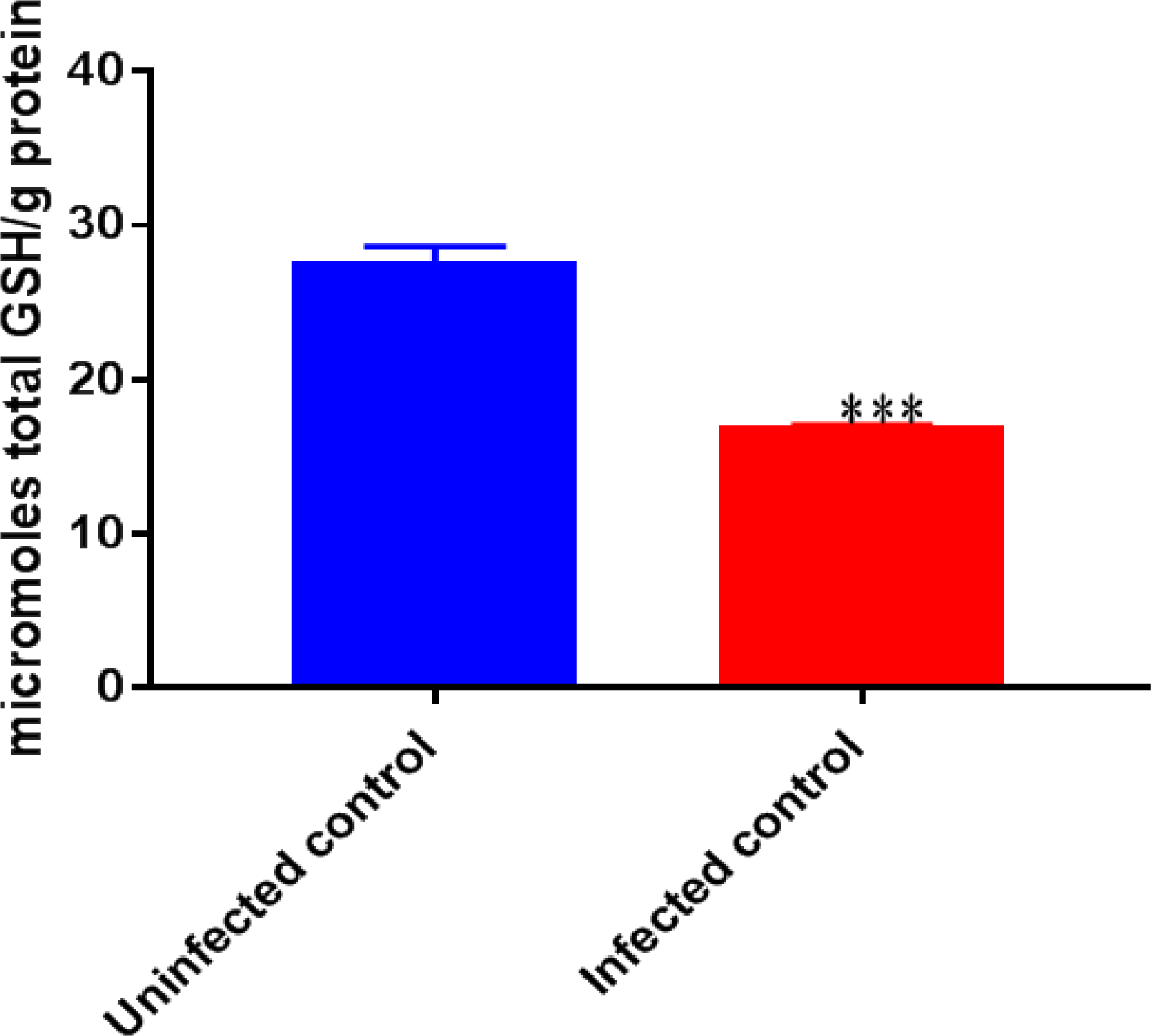
Levels of total GSH in uninfected and M. *tb*-infected THP-1 cells. GSH assay was performed by colorimetric method using am assay kit from Arbor Assays. Corrections were made to total protein measured by BCA Protein Assay Kit from Thermo Scientific. GSH and total protein levels were measured from the cellular lysates. There was a notable decrease in the levels of GSH when THP-1 cells were infected with the Erdman strain of M. *tb*. Data represent means ±SE from 6 trials. ***p<0.0005 when comparing infected cells with uninfected.

### Survival of Erdman strain of *M*. *tb* inside untreated THP-1 macrophages

There was a statistically significant increase in the intracellular survival of *M. tb* residing inside the untreated macrophages between the initial (1 hour) and final time point of termination (12-days post-infection) [Fig. 1B]. These results signify the ability of Erdman strain of *M*. *tb* to successfully survive and replicate inside the THP-1 cells.

**Fig. 1B:**
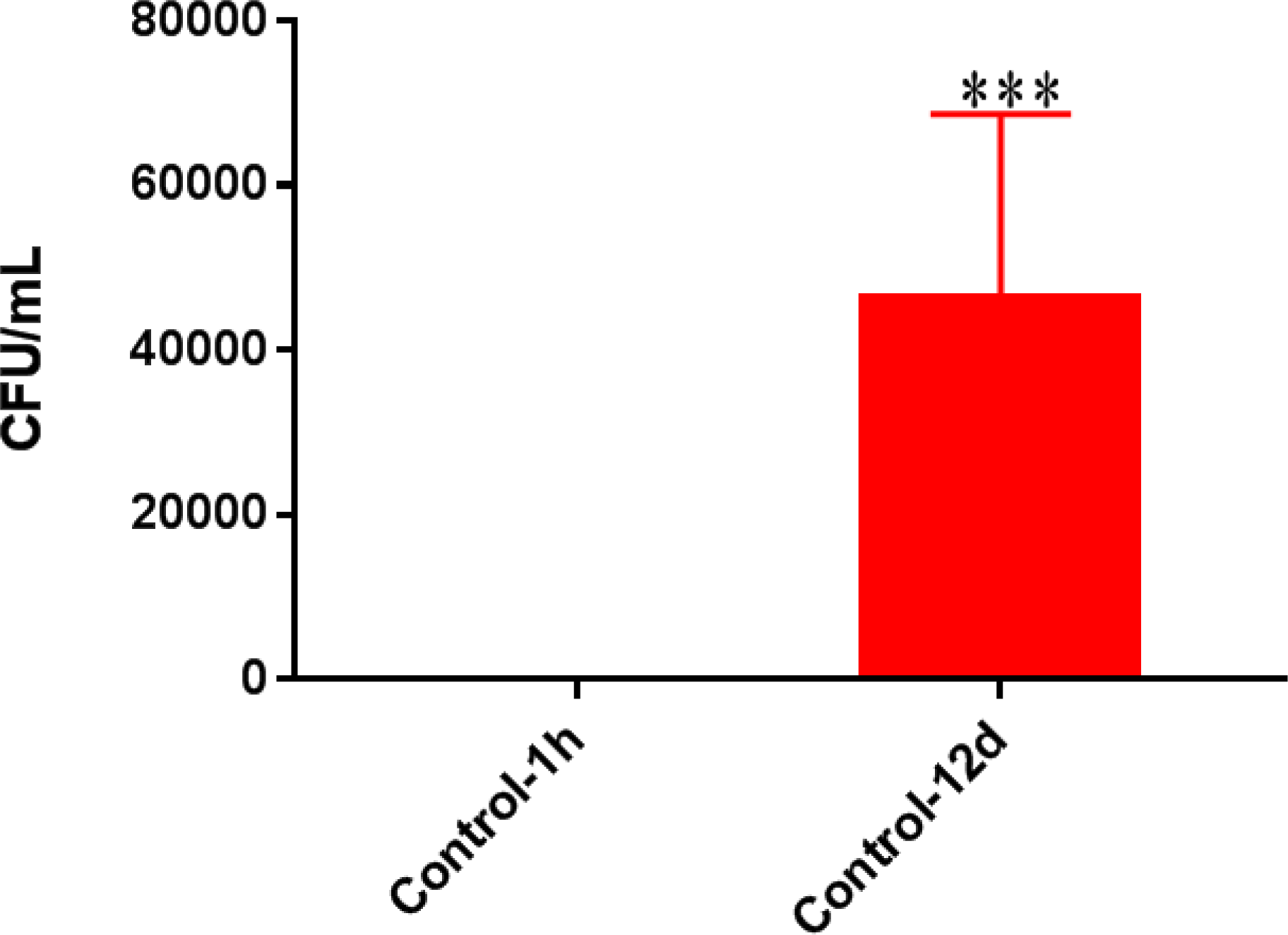
Survival of Erdman strain of *M*. *tb* inside untreated THP-1 macrophages. THP-1 cells were cultured in RPMI containing 10% FBS, and allowed to differentiate into macrophages by the addition of PMA at a concentration of 10 ng/ml. There was a significant increase in bacterial numbers when THP-1 cells were infected with M. *tb*. Data represent means ±SE from 6 trials. ***p<0.0005 when comparing 1 hour with 12-day time point of termination.

### Levels of TNF-α in the supernatants of uninfected and *M*. *tb*-infected THP-1 cells

Levels of TNF-α, a pro-inflammatory cytokine were quantified in the supernatants collected from the uninfected macrophages and *M. tb*-infected macrophages by “sandwich” ELISA. When compared to the uninfected control category, there was a statistically significant six-fold increase in the levels of TNF-α in *M*. *tb*-infected macrophages (Fig. 1C). Our results imply that excess production of TNF-α by *M*. *tb*-infected macrophages result in GSH decrease thereby favoring the intracellular survival of *M*. *tb.*

**Fig. 1C:**
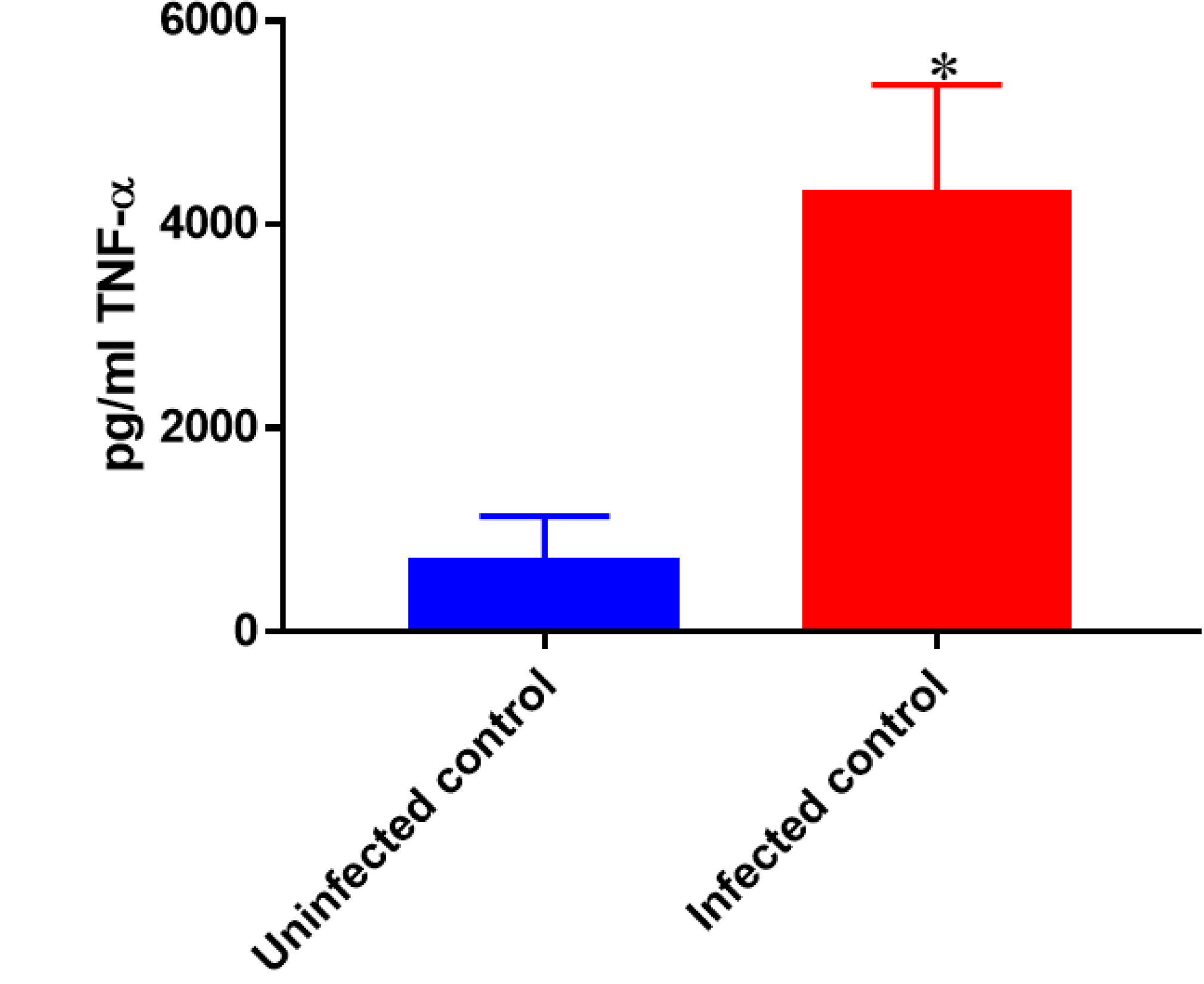
Levels of TNF-α in the supernatants of uninfected and *M*. *tb*-infected THP-1 cells. Assay of TNF-α was performed using an ELISA Ready-Set-Go kit from eBioscience. There was a significant increase in the levels of TNF-α when macrophages were infected with M. *tb*. Data represent means ±SE from 6 trials. *p<0.05 when comparing infected samples to uninfected controls.

### Measurement of GSH, bacterial survival, TNF-α and IL-10 levels in *M. tb*-infected and M. *tb*-infected + NAC-treated THP-1 cells

Treatment of *M*. *tb*-infected macrophages with NAC resulted in a significant two-fold increase in the intracellular levels of GSH when compared to the *M*. *tb* infected sham control group (Fig. 2A). NAC-treatment therefore restores the levels of GSH in *M*. *tb*-infected macrophages. Restoration in the levels of GSH correlated with significant reduction in the intracellular survival of *M*. *tb* inside the NAC-treated macrophages (Fig. 2B). GSH-replenishment in *M*. *tb*-infected macrophages also resulted in a significant 8-fold decrease in levels of TNF-α compared to the *M*. *tb*-infected control group (Fig. 2C). Importantly, NAC-treatment of *M*. *tb*-infected macrophages resulted in a 4-fold decrease in levels of IL-10 compared to the *M*. *tb*-infected sham control group (Fig. 2D). These results further confirm that by restoring redox homeostasis and cytokine balance there is improved control of intracellular *M*. *tb* infection.

**Fig. 2A:**
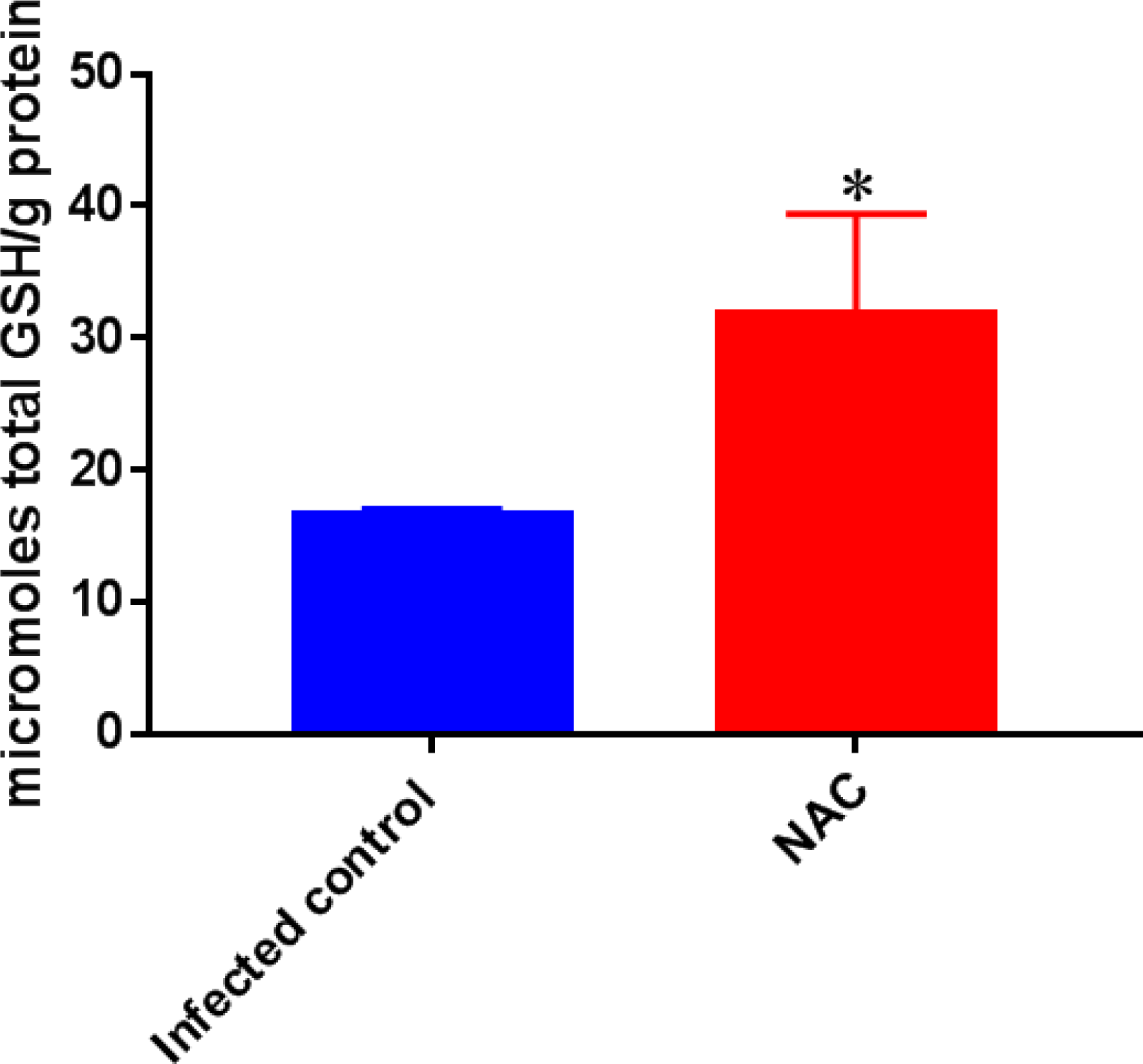
Levels of total GSH in *M*. *tb*-infected and *M*. *tb*-infected + NAC-treated THP-1 cells. GSH assay was performed by colorimetric assay using an assay kit from Arbor Assays. Corrections were made to total protein measured by BCA Protein Assay Kit from Thermo Scientific. GSH and total protein levels were measured from cellular lysates. There was a significant increase in the levels of GSH when *M. tb*-infected samples were treated with NAC. Data represent means ±SE from 6 trials. *p<0.05 when comparing *M*. *tb*-infected samples treated with NAC to infected sham controls.

**Fig. 2B:**
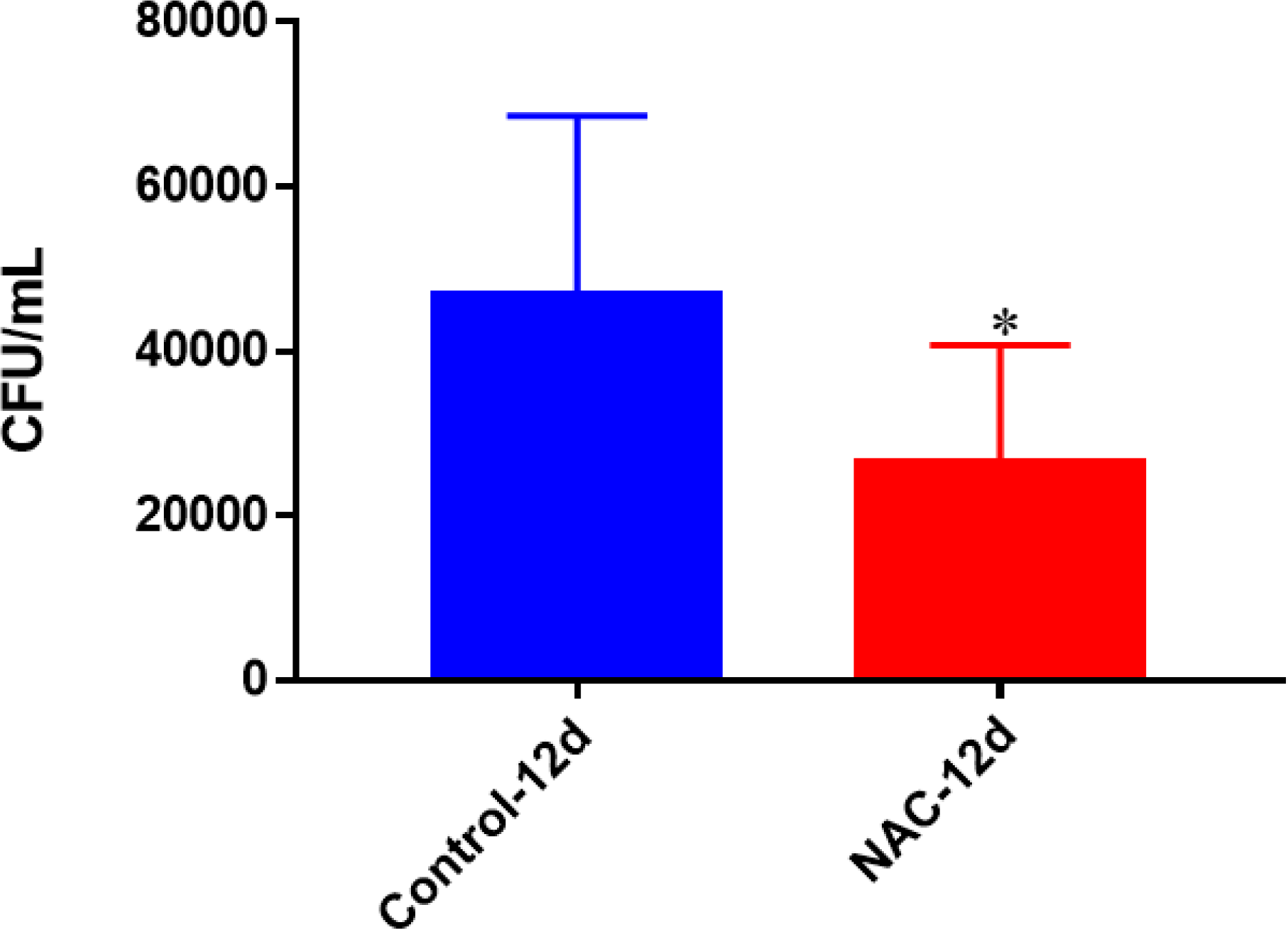
Survival of Erdman strain of *M. tb* inside untreated and NAC-treated THP-1 macrophages. THP-1 cells were cultured in a medium of RPMI and 10% FBS, and allowed to differentiate into macrophages by addition of PMA at a concentration of 10 ng/ml. There was a significant decrease in bacterial numbers when *M. tb*-infected macrophages were treated with NAC. Data represent means ±SE from 6 trials. *p<0.05 when comparing *M*. *tb*-infected macrophages treated with NAC to infected sham controls.

**Fig. 2C:**
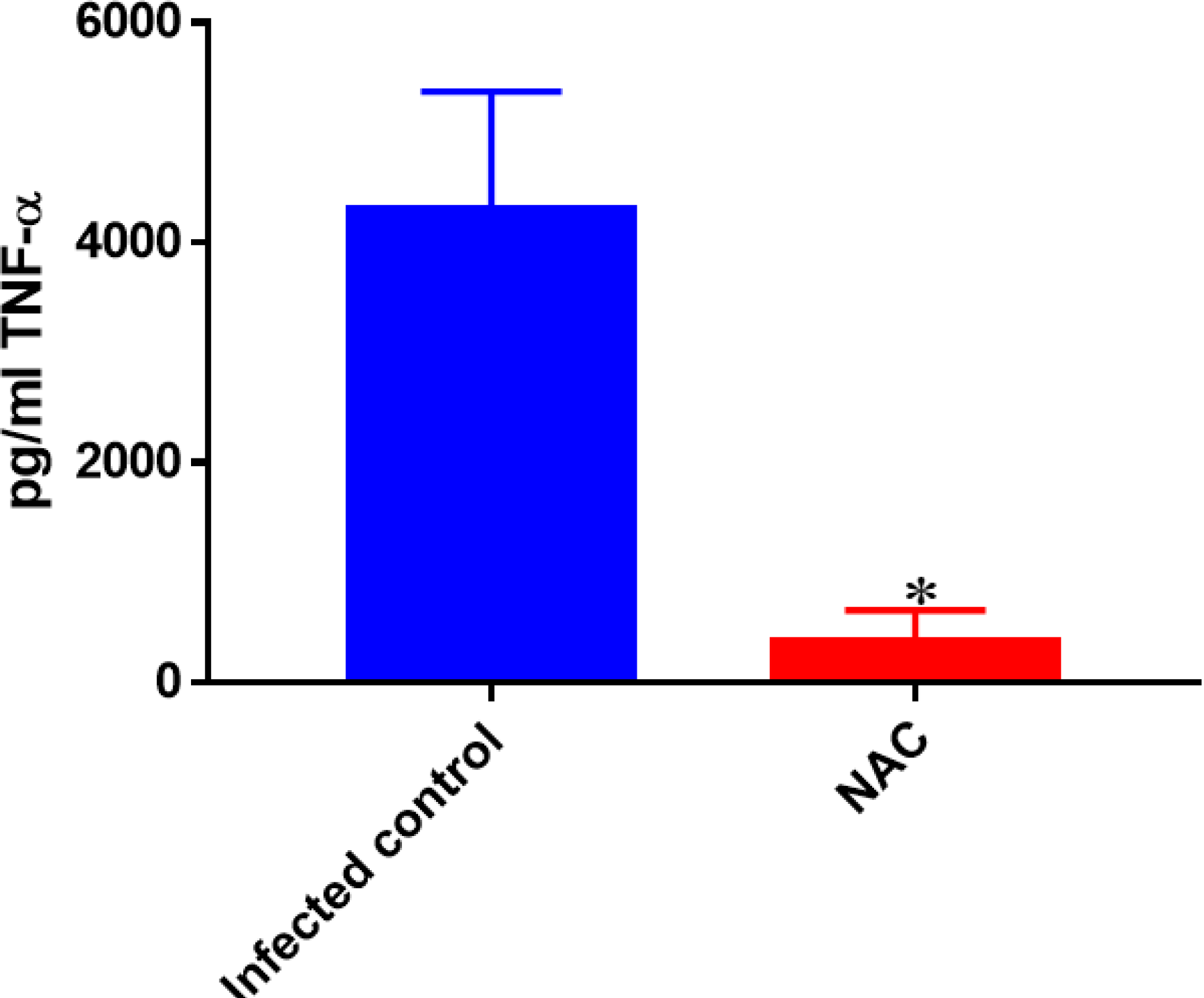
Assay of TNF-α in the supernatants from *M*. *tb*-infected and *M*. *tb*-infected + NAC-treated THP-1 cells. Assay of TNF-α was performed using an ELISA Ready-Set-Go kit from eBioscience. There was a significant decrease in the levels of TNF-a when macrophages were infected with *M. tb* and treated with NAC. Data represent means ±SE from 6 trials. *p<0.05 when comparing *M. tb*-infected macrophages treated with NAC to infected sham controls.

**Fig. 2D:**
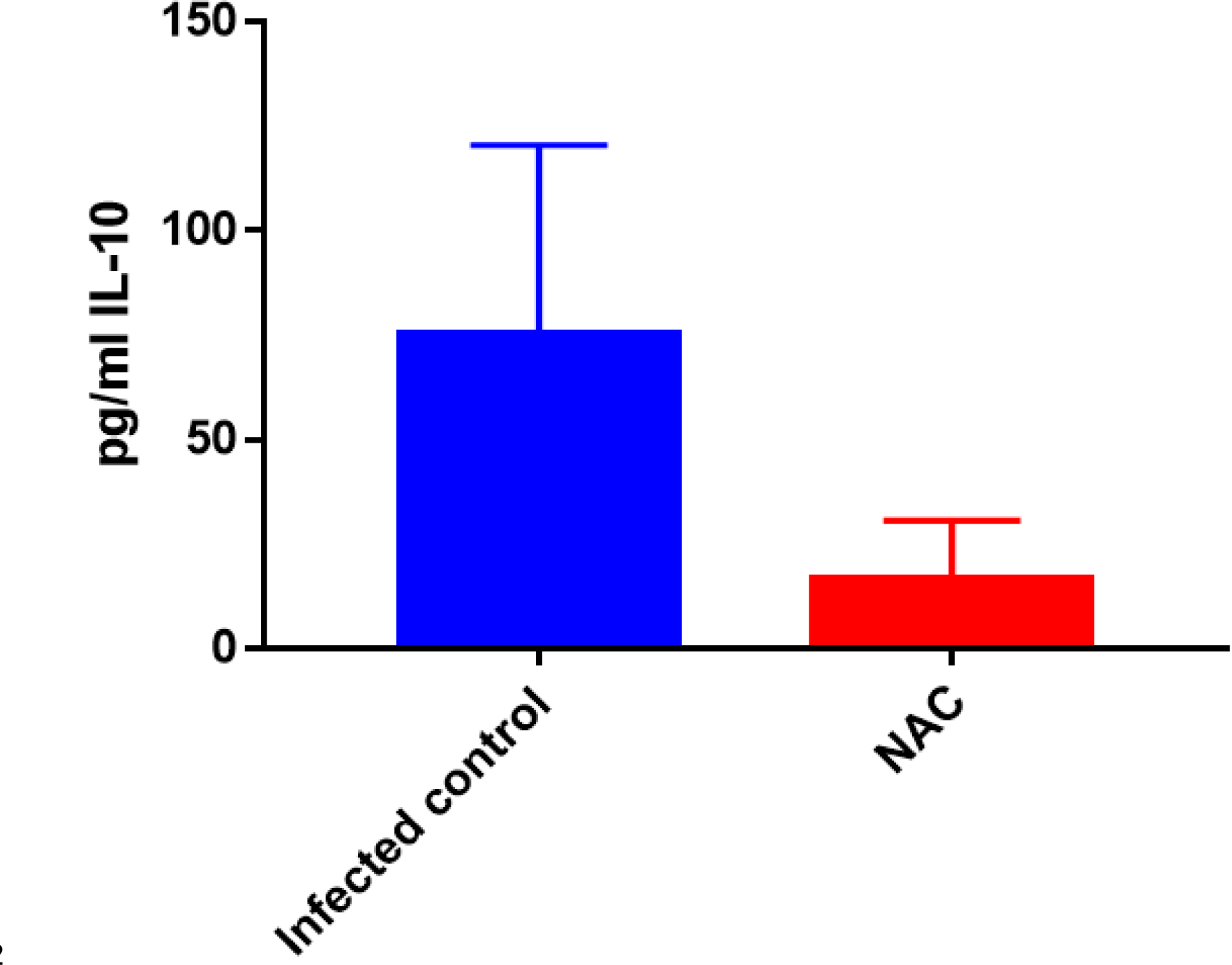
Assay of IL-10 in the supernatants from *M*. *tb*-infected and *M. tb*-infected + NAC-treated THP-1 cells. IL-10 was measured using an ELISA Ready-Set-Go kit from eBioscience. Although there was a decrease in the levels of IL-10 when M. *tb*-infected macrophages were treated with NAC, this difference was not found to be statistically significant. Data represent means ±SE from 6 trials.

### Quantification of GSH levels, *M. tb* survival, TNF-α and IL-10 levels in *M. tb*-infected, M. *tb*-infected + INH-treated and *M*. *tb*-infected with INH + NAC-treated THP-1 cells

INH is the first line antibiotic most commonly used for treatment of *M. tb* infection. *M*. *tb*-infected macrophages were treated as follows: sham-treatment, treated with INH (0.125 micrograms/mL), and treated with combination of INH (0.125 micrograms/mL) +NAC (10mM), and the effects of INH and INH+NAC treatments in altering the levels of GSH and cytokines, and *M. tb* survival was determined. When compared to the infected sham control, there was a significant increase in the levels of GSH in *M. tb*-infected macrophages treated with INH and INH+ NAC (Fig. 3A). In comparison to INH alone category, treatment with INH+NAC resulted in complete clearance of M. *tb* infection (Fig. 3B). Treatment with either INH or the combination of INH+ NAC resulted in almost 10,000-fold decrease in the viability of *M*. *tb* when compared to treatment with NAC only (Fig 2B), and roughly 15,000-fold decrease in the viability of *M*. *tb* compared to sham control group (Fig. 1B). These results further emphasize that reduction in *M. tb* burden can restore the intracellular levels of GSH. Treatment of *M*. *tb*-infected macrophages with INH resulted in a statistically significant decrease in the levels of TNF-α when compared to the infected sham control group. Treatment of *M*. *tb*-infected macrophages with INH + NAC also resulted in a statistically significant decrease in the levels of TNF-α (Fig. 3C). INH + NAC treatment resulted in further reduction in the levels of TNF-α in comparison to treatment with INH alone. Consistent with our findings from the TNF-α assay, INH treatment of *M. tb*-infected macrophages also resulted in reduction in the levels of IL-10 when compared to the infected sham controls. INH + NAC treatment caused additional decrease in the levels of IL-10 (Fig. 3D). These findings specify the effects of *M*. *tb* clearance in diminishing the levels of TNF-α and IL-10.

**Fig. 3A:**
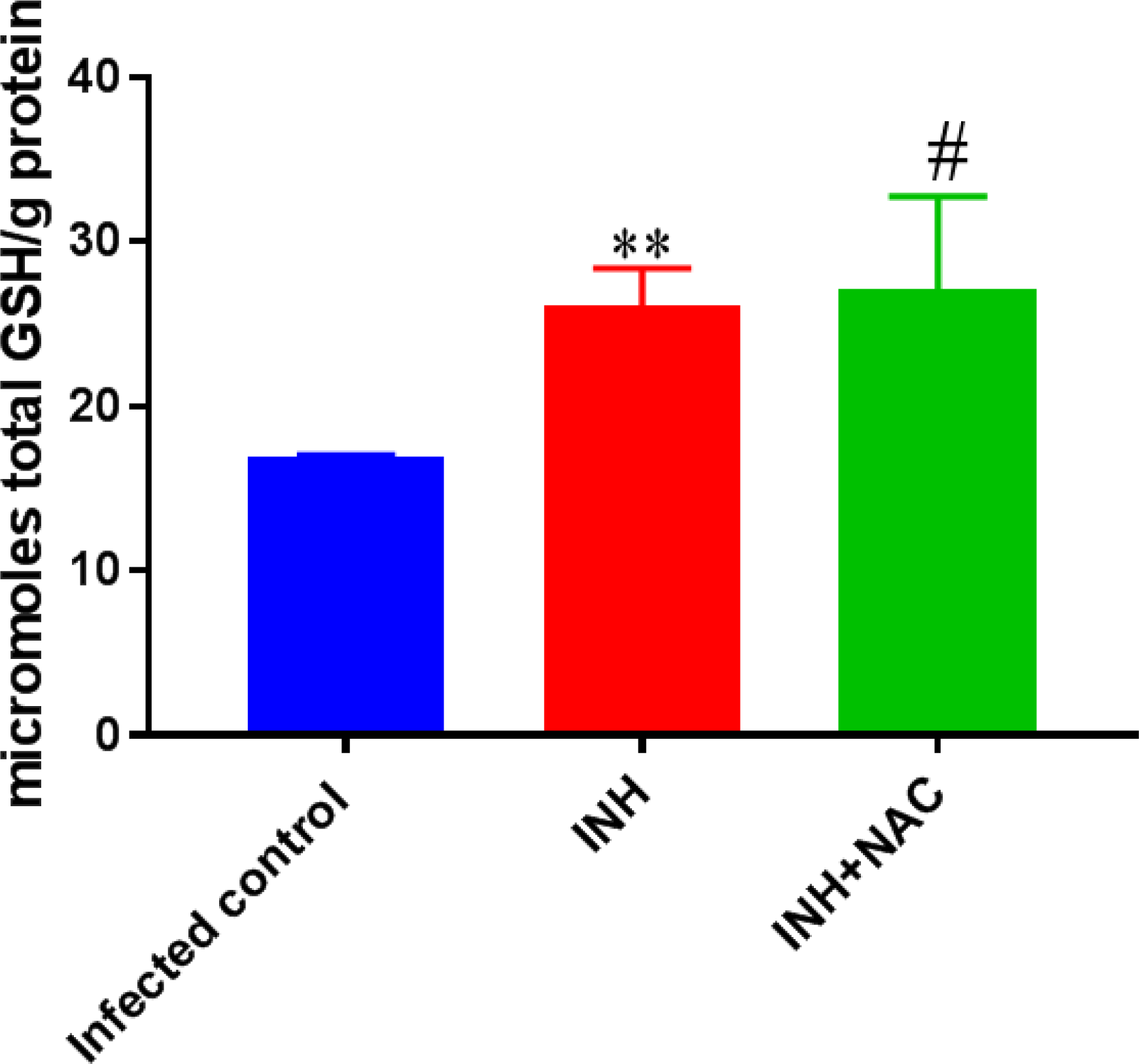
Levels of total GSH in M. *tb*-infected, *M*. *tb*-infected + INH-treated and *M*. *tb*-infected + INH + NAC-treated THP-1 cells. GSH assay was performed using a colorimetric assay kit from Arbor Assays. There was a significant increase in the levels of GSH when M. *tb*-infected macrophages were treated with INH and INH+NAC. Data represent means ±SE from 6 trials. **p<0.005 when comparing infected samples treated with INH to infected sham controls. ^#^p<0.05 when comparing infected samples treated with INH+NAC to infected sham controls.

**Fig. 3B:**
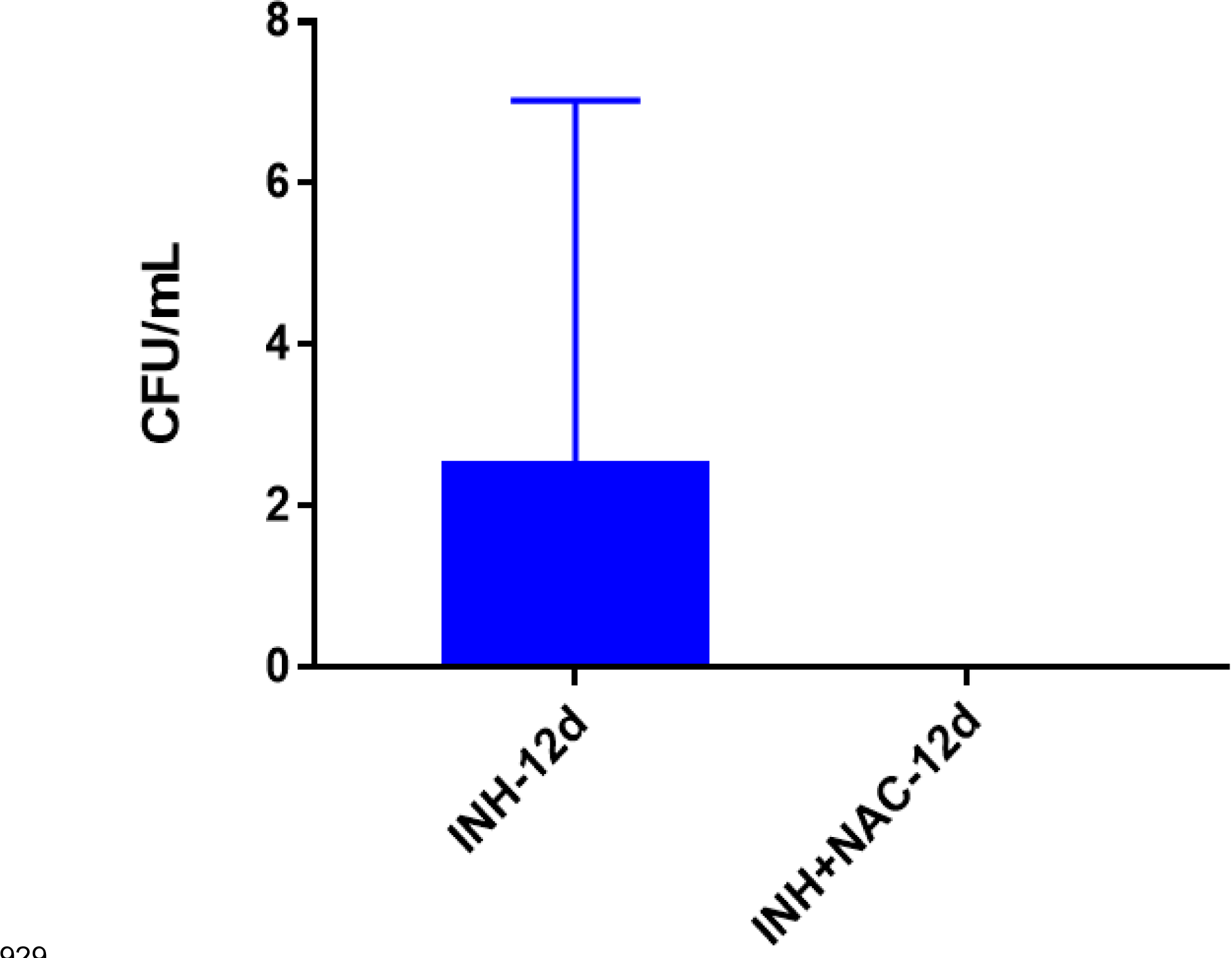
Survival of Erdman strain of *M*. *tb* inside INH and INH + NAC-treated THP-1 macrophages. THP-1 cells were cultured in a medium of RPMI and 10% FBS and allowed to differentiate into macrophages by addition of PMA at a concentration of 10 ng/ml. INH+NAC treatment resulted in clearance of *M*. *tb* infection compared to treatment with INH only. Data represent means ±SE from 6 trials.

**Fig. 3C:**
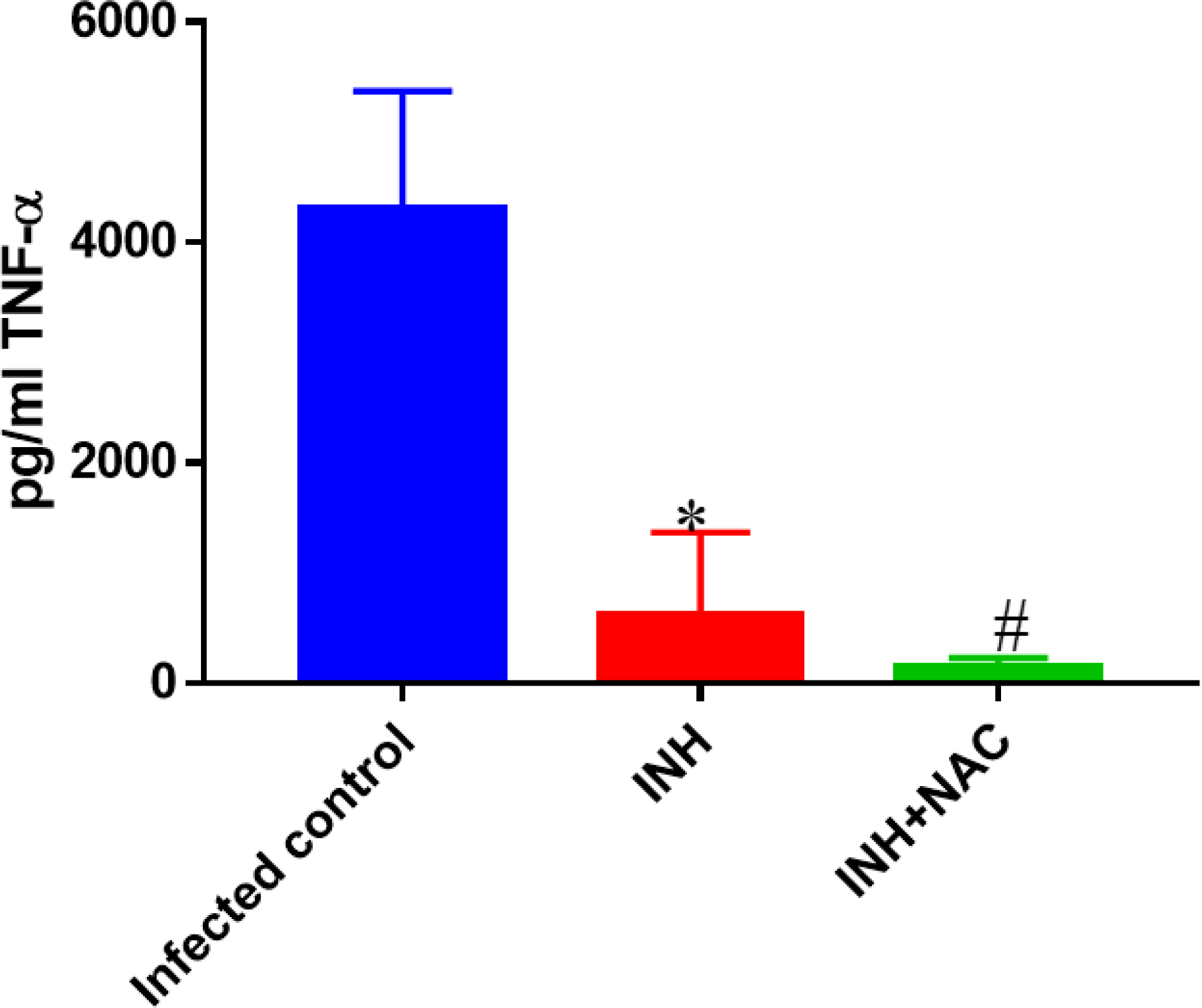
Assay of TNF-α in the supernatants from INH and INH + NAC-treated THP-1 cells. Assay of TNF-α was performed using an ELISA Ready-Set-Go kit from eBioscience. There was a significant decrease in the levels of TNF-α when samples were infected with *M*. *tb* and treated with either INH or INH+NAC. Data represent means ±SE from 6 trials. *p<0.05 when comparing M. *tb*-infected macrophages treated with INH to infected sham controls. ^#^p<0.05 when comparing M. *tb*-infected macrophages treated with INH+NAC to infected sham controls.

**Fig. 3D:**
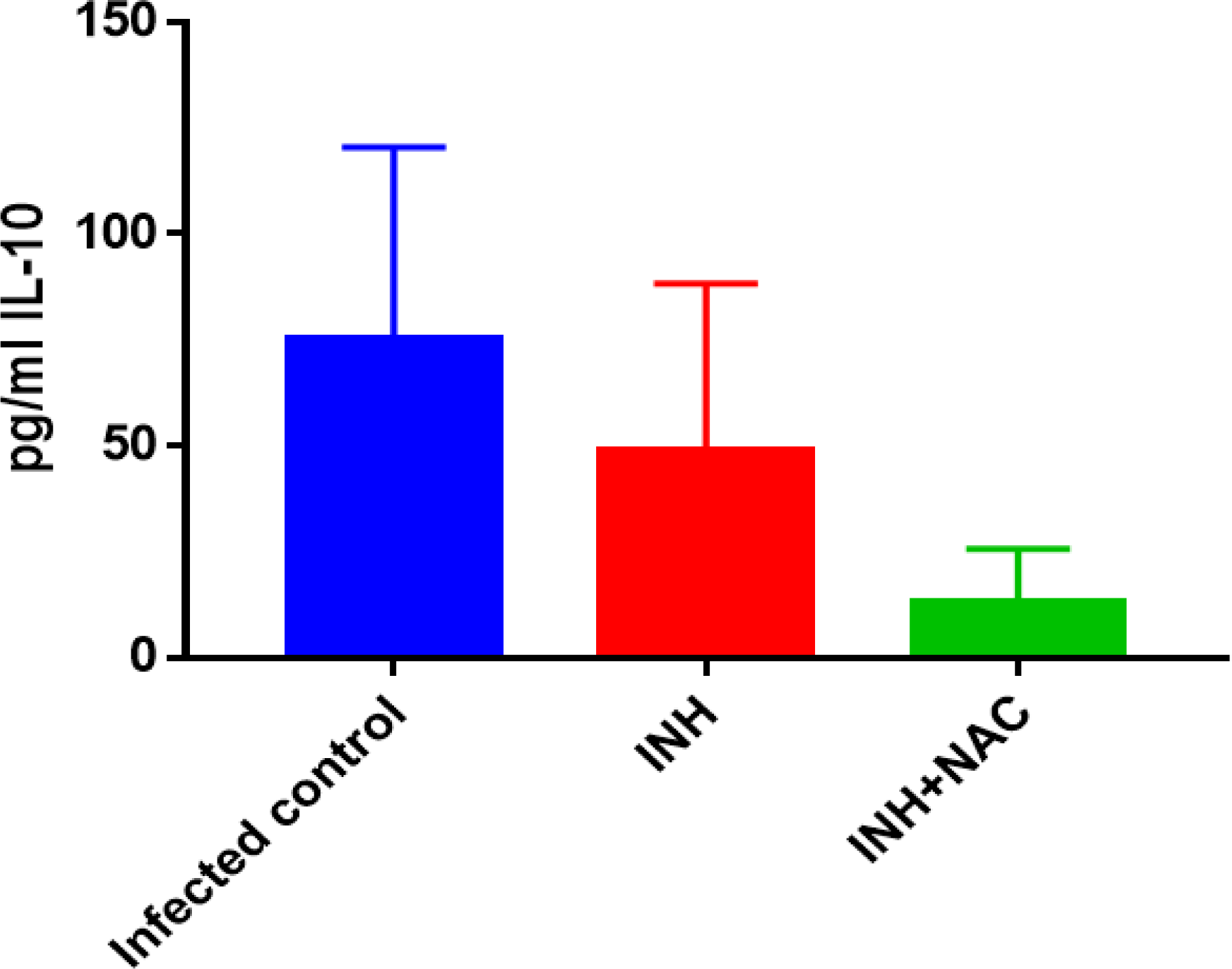
Assay of IL-10 in the supernatants from INH and INH + NAC-treated THP-1 cells. IL-10 levels were measured using an ELISA Ready-Set-Go kit from eBioscience. Although there was a decrease in the levels of IL-10 when samples were infected with *M*. *tb* and treated with INH or INH+NAC, this difference was not found to be statistically significant. Data represent means ±SE from 6 trials.

### Quantification of GSH levels, *M. tb* survival, TNF-α and IL-10 levels in *M*. *tb*-infected, *M*. *tb*-infected + RIF-treated and *M*. *tb*-infected with RIF+ NAC-treated THP-1 cells

RIF is another dominant first-line antibiotic used for the treatment of TB. GSH levels were therefore measured in the *M*. *tb-*infected sham control macrophages and *M*. *tb*-infected macrophages treated with either RIF alone (0.125 micrograms/ml) or RIF (0.125 micrograms/ml) in combination with NAC (10mM). The RIF-treatment to infected macrophages resulted in a significant increase in GSH compared to the infected sham control group. Importantly, RIF + NAC treatment resulted in further enhancement in the levels of GSH and a statistically significant increase compared to both infected sham control category and also RIF unaided category (Fig. 4A).

**Fig. 4A:**
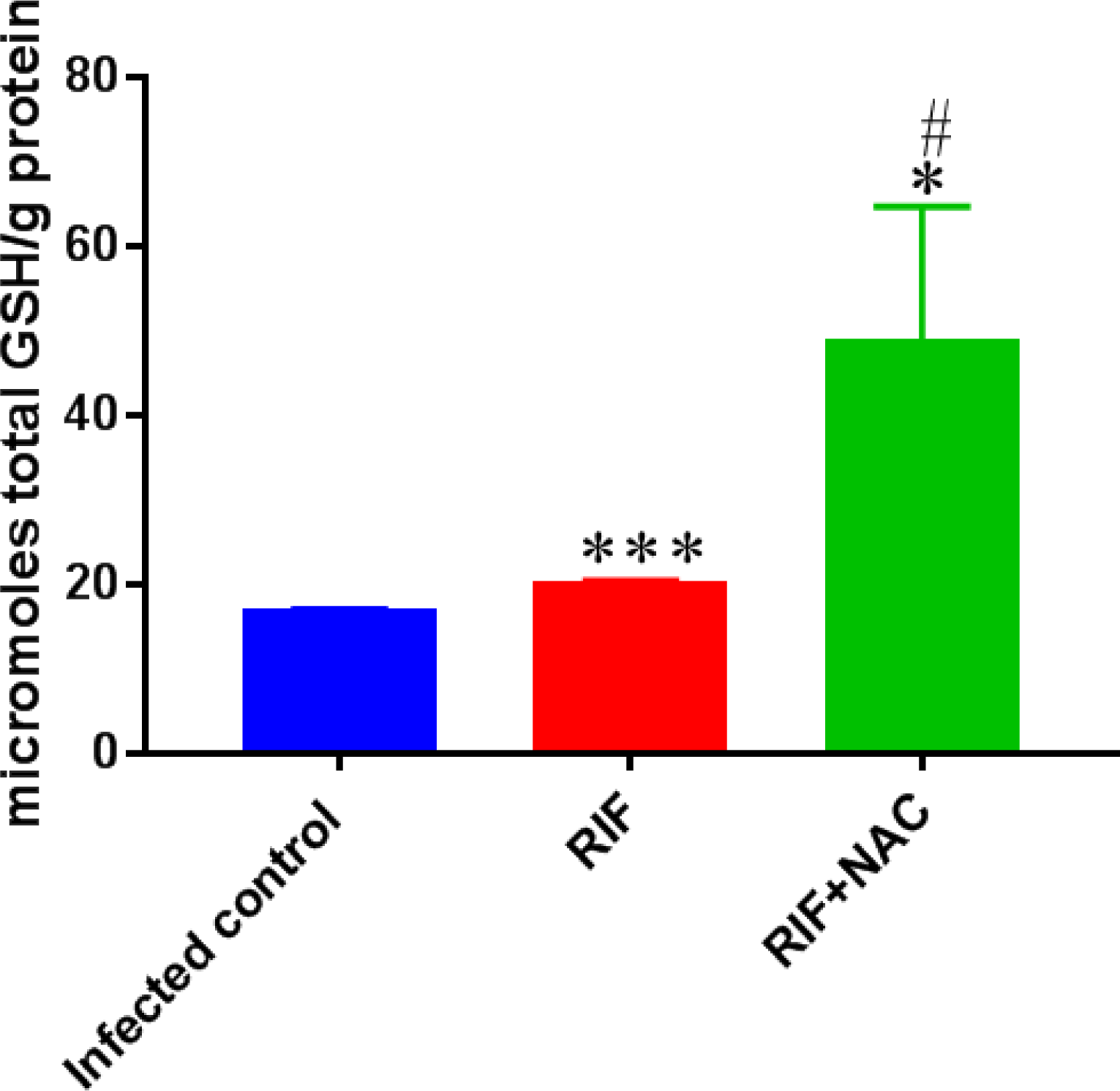
Levels of total GSH in M. *tb*-infected, M. *tb*-infected + RIF-treated and *M*. *tb*-infected + RIF + NAC-treated THP-1 cells. GSH assay was performed using a colorimetric assay kit from Arbor Assays. There was a significant increase in the levels of GSH when *M*. *tb*-infected macrophages were treated with RIF and RIF+NAC. Data represent means ±SE from 6 trials. ***p<0.0005 when comparing infected macrophages treated with RIF to infected sham controls. ^#^p<0.05 when comparing infected macrophages treated with RIF+NAC to RIF only. *p<0.05 when comparing infected macrophages treated with RIF+NAC to infected sham controls.

The notable increase in the levels of GSH in RIF + NAC treatment group was accompanied by significant and 8-fold reduction in the intracellular viability of *M. tb* compared to treatment with RIF alone (Fig. 4B). When comparing the efficacy of INH versus RIF, there was also approximately 70 times more intracellular *M. tb* when macrophages were treated with RIF instead of INH, and 25 times more bacteria when treatment of RIF with NAC was used instead of INH with NAC (Fig. 3B). When compared to sham treatment, RIF treatment reduced bacterial burden by almost 200 times (Fig. 1B) and treatment of RIF+NAC reduced the *M. tb* counts by 1,600 times. Treatment of *M. tb*-infected macrophages with combination of RIF+ NAC, also resulted in a statistically significant decrease in the levels of TNF-α compared to both the infected sham control and lone RIF (Fig. 4C). Treatment with RIF caused 50% reduction in the levels of TNF-α compared to sham-treated control (Fig. 4C). In comparison to sham-control, treatment with RIF+NAC resulted in 80-fold reduction in the levels of TNF-α (Fig. 4C). RIF administered alone was not able to lower the levels of TNF-α as much as INH by itself, but the addition of NAC to RIF, lowered the TNF-α levels 8 times more than that of INH and NAC treatment (Fig. 3C). RIF treatment resulted in a decrease in the levels of IL-10 compared to the sham-control group (Fig 4D). Treatment of M. *tb*-infected macrophages with combination of RIF+ NAC, resulted in a further decrease in the levels of IL-10 compared to the control and lone RIF, but neither of these decreases were statistically significant (Fig 4D). IL-10 trends were very similar when comparing RIF and INH treatments. However, INH was twice as effective in lowering the levels of IL-10 in macrophage cultures (Fig. 3D).

**Fig. 4B:**
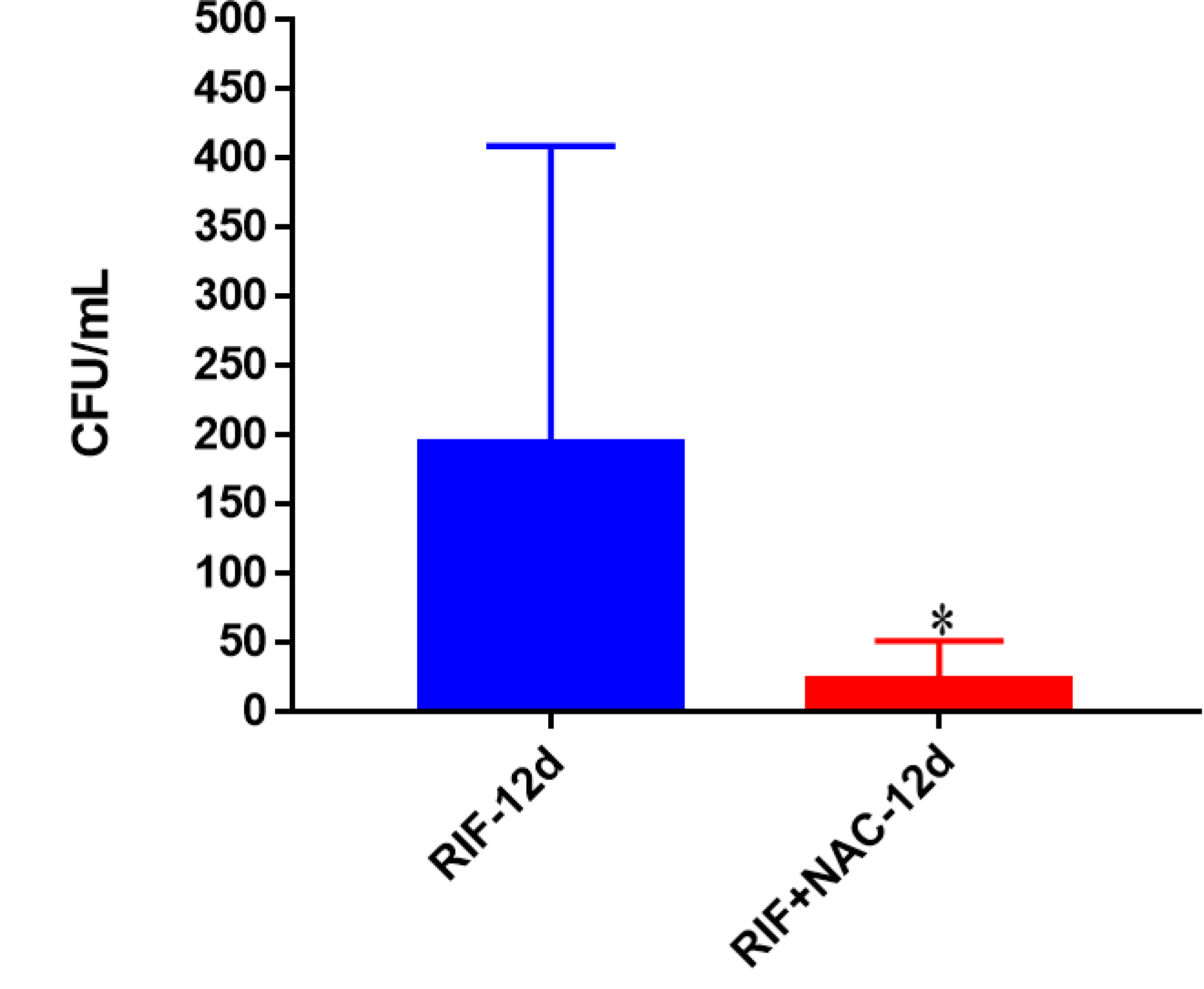
Survival of Erdman strain of *M. tb* inside RIF and RIF + NAC-treated THP-1 macrophages. THP-1 cells were cultured in a medium of RPMI and 10% FBS, and allowed to differentiate into macrophages by addition of PMA at a concentration of 10 ng/ml. There was a significant reduction in the bacterial numbers when THP-1 cells were treated with RIF+NAC compared to RIF only. Data represent means ±SE from 6 trials. *p<0.05 when comparing infected samples treated with RIF+NAC to RIF only treatment.

**Fig. 4C:**
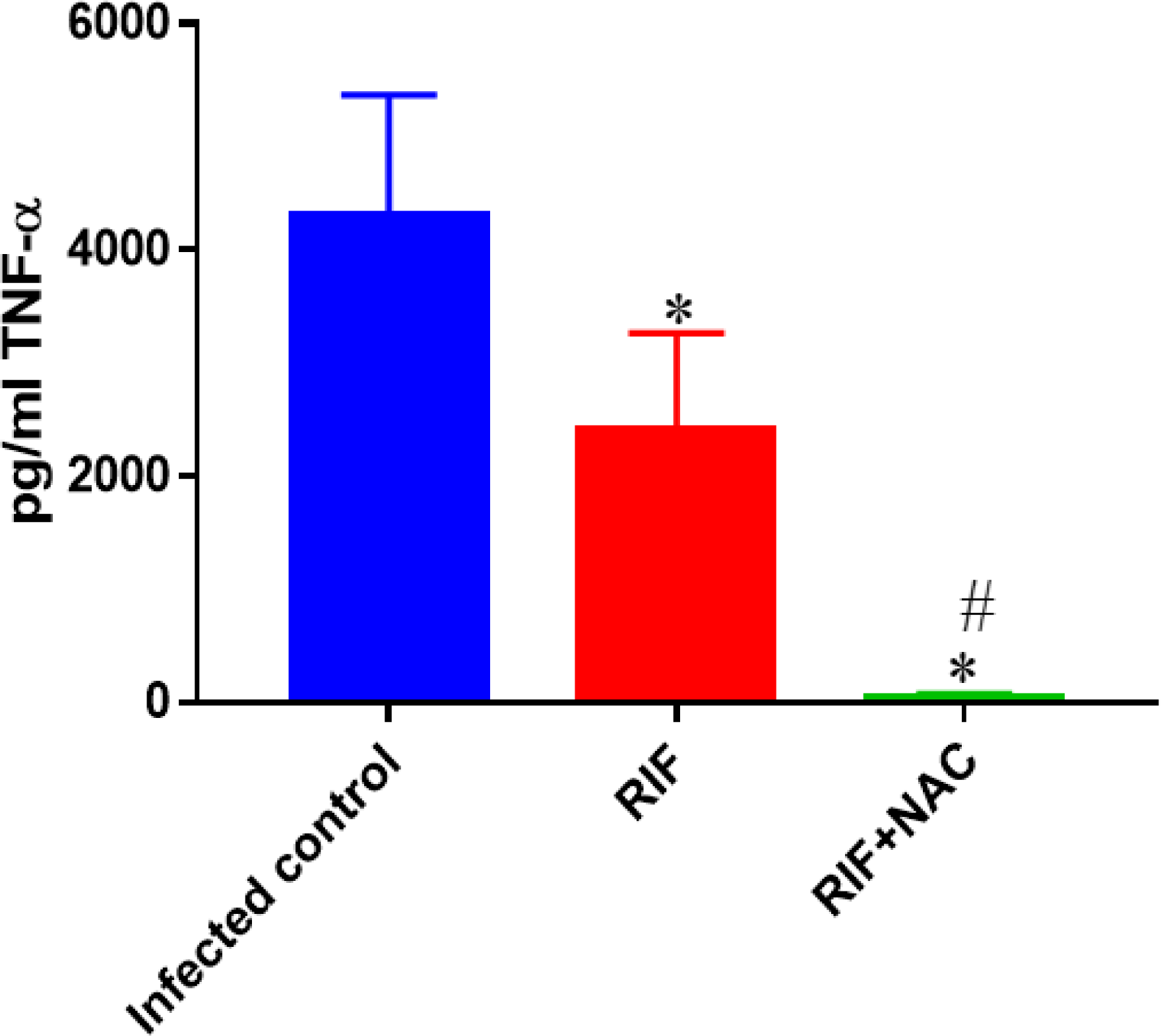
Assay of TNF-α in the supernatants from RIF and RIF + NAC-treated THP-1 cells. Assay of TNF-α was performed using an ELISA Ready-Set-Go kit from eBioscience. There was a significant decrease in levels if TNF-α when macrophages were infected with *M*. *tb* and treated with RIF or RIF+NAC. Data represent means ±SE from 6 trials. *p<0.05 when comparing infected macrophages treated with RIF or RIF+NAC to infected controls. ^#^p<0.05 when comparing infected macrophages treated with RIF+NAC to treatment with RIF only.

**Fig. 4D:**
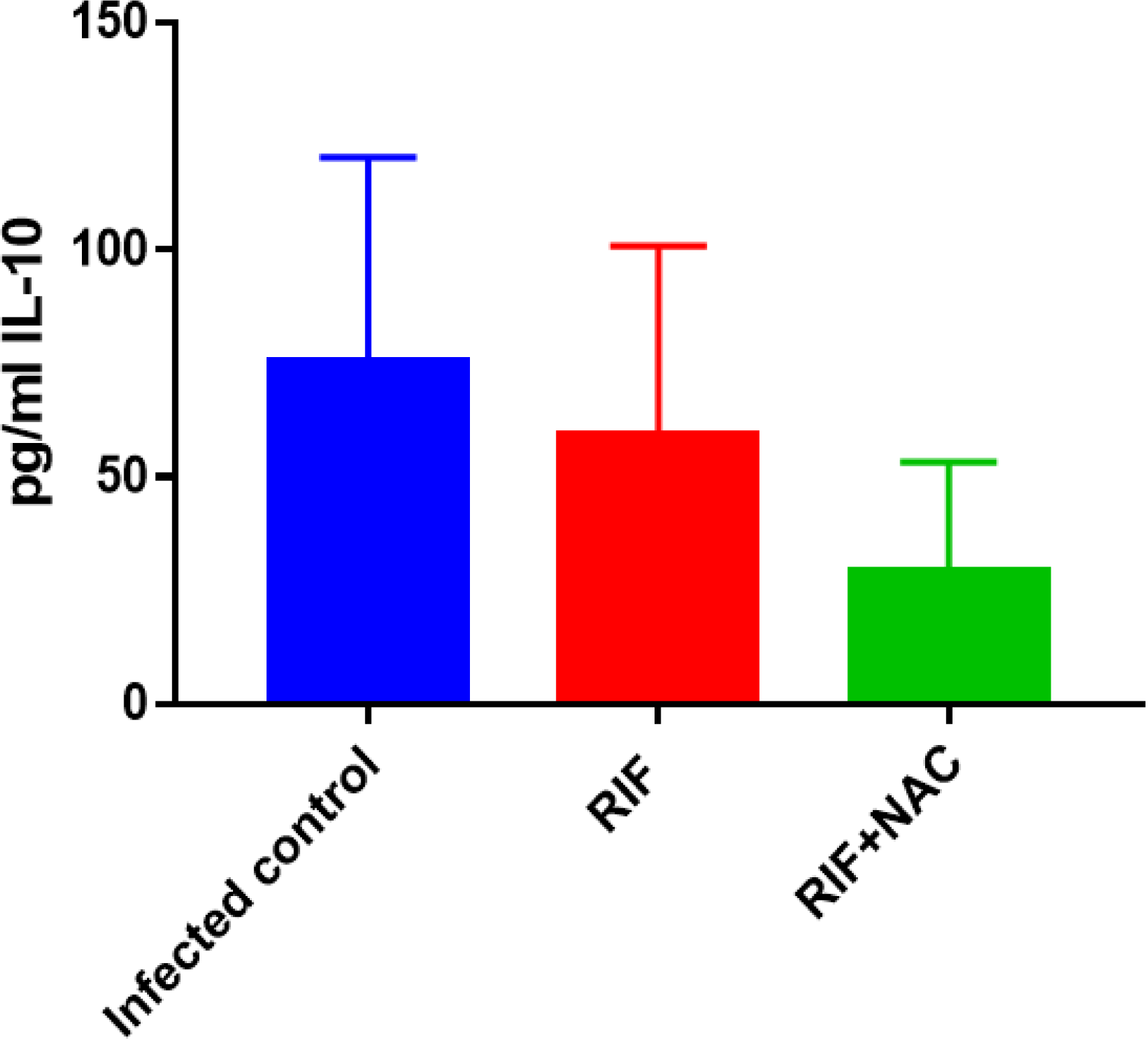
Assay of IL-10 in the supernatants from RIF and RIF + NAC-treated THP-1 cells. IL-10 was measured using an ELISA Ready-Set-Go kit from eBioscience. Although there was a decrease in levels of IL-10 when macrophages were infected with *M. tb* and treated with RIF or RIF+NAC, this difference was not found to be statistically significant. Data represent means ±SE from 6 trials.

### Quantification of GSH levels, *M*. *tb* survival, TNF-α and IL-10 levels in *M*. *tb*-infected, *M. tb*-infected + EMB-treated and *M*. *tb*-infected with EMB+ NAC-treated THP-1 cells

EMB is an antibiotic that is normally given with a myriad of other first line antibiotics for TB treatment. We tested the efficacy of stand-alone EMB (8.0 micrograms/ml), as well as the efficacy of EMB (8.0 micrograms/ml) supplemented with NAC (10mM) in altering the levels of GSH and cytokines, and *M*. *tb* survival inside the macrophages. GSH levels were increased by 2-fold in the macrophages treated with EMB compared to the infected sham control group, although this increase was not significant. However, when macrophages were treated with EMB+NAC, there was a statistically significant 3-fold increase in the levels of GSH compared to the sham control category (Fig. 5A). Consistent with the increase in the levels of GSH, there was four-fold decrease in the intracellular survival of *M*. *tb* inside the EMB+NAC-treated macrophages compared to the stand-alone EMB treated macrophages (Fig. 5B). When compared to the infected sham control group, the infected macrophages treated with stand-alone EMB had significantly reduced levels of TNF-α (Fig. 5C). EMB+NAC treatment resulted in further decrease in the levels of TNF-α in comparison to sham-control and EMB-alone groups. EMB and EMB+NAC treatments resulted in more than 50% and 80% decrease respectively, in the levels of TNF-α in comparison to sham-control group (Fig. 5C). Although not significant, a distinct reduction in the levels of IL-10 was observed when *M*. *tb* infected macrophages were treated with EMB+ NAC (Fig. 5D).

**Fig. 5A:**
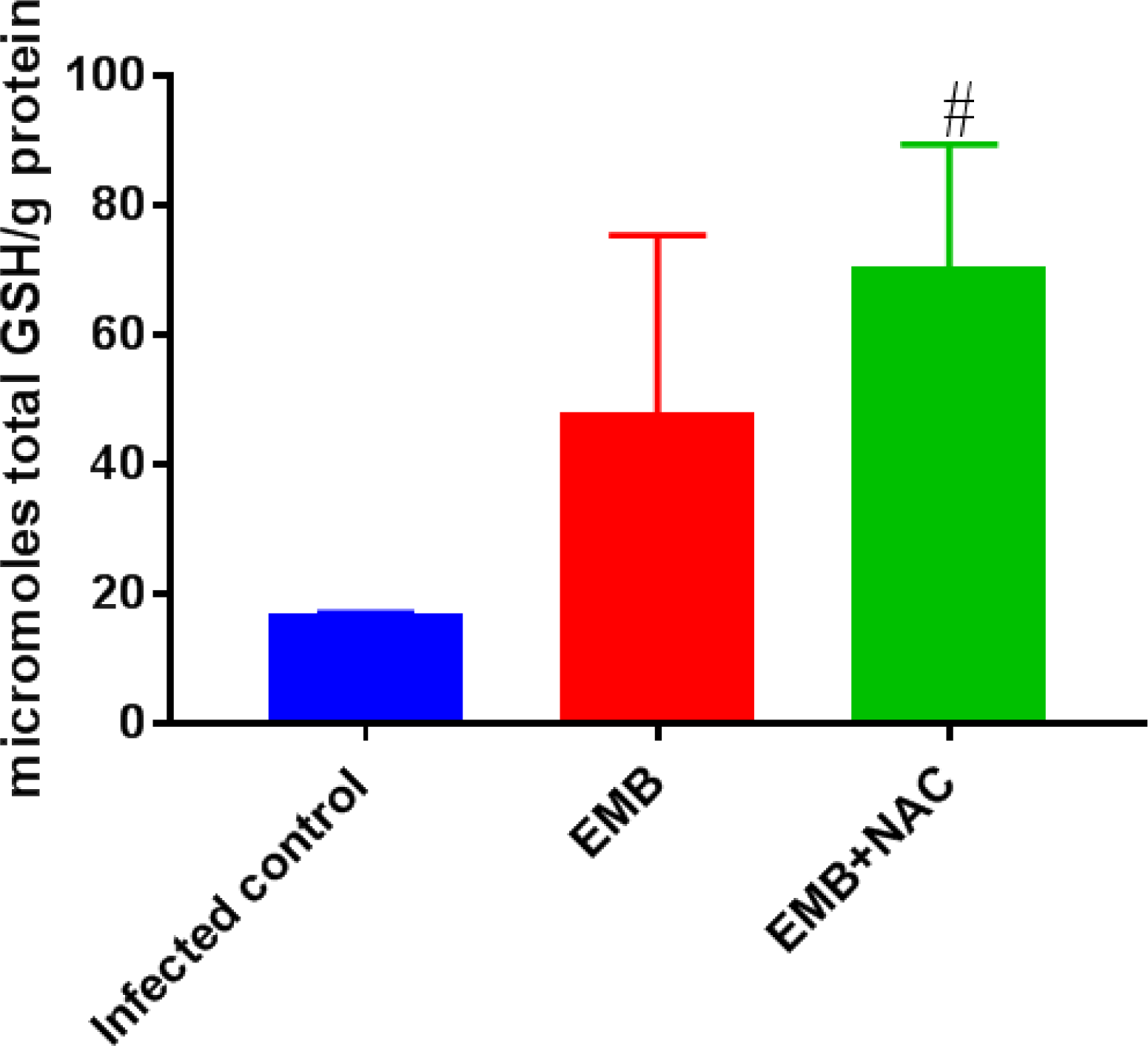
Levels of total GSH in *M. tb*-infected, *M*. *tb*-infected + EMB-treated and *M. tb*-infected + EMB + NAC-treated THP-1 cells. GSH assay was performed using a colorimetric assay kit from Arbor Assays. There was a significant increase in the levels of GSH when infected macrophages were treated with EMB+NAC. Data represent means ±SE from 6 trials. ^#^p<0.05 when comparing infected samples treated with EMB+NAC to infected controls.

**Fig. 5B:**
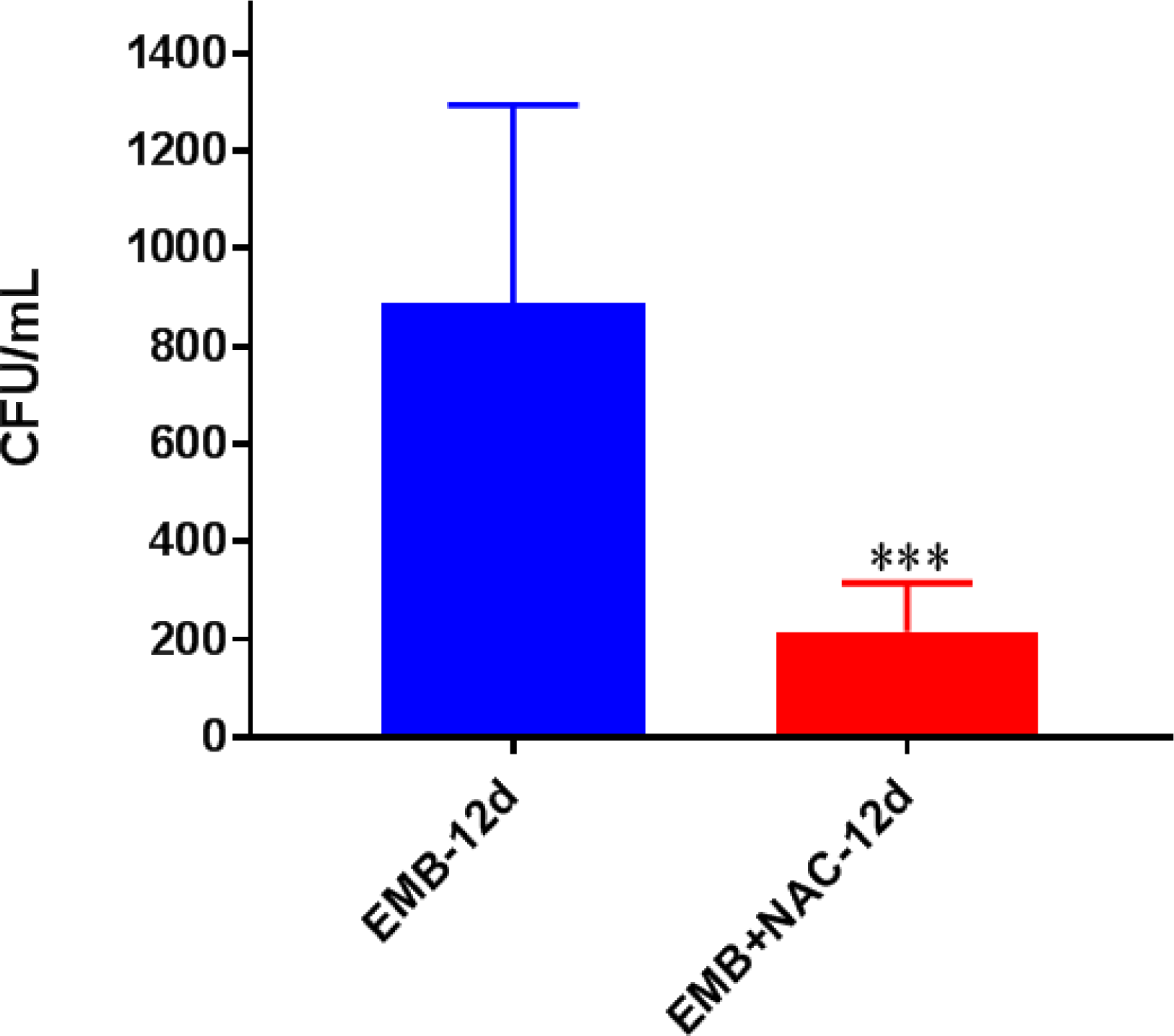
Survival of Erdman strain of *M*. *tb* inside EMB and EMB + NAC-treated THP-1 macrophages. THP-1 cells were cultured in a medium of RPMI and 10% FBS, and allowed to differentiate into macrophages by addition of PMA at a concentration of 10 ng/ml. There was a significant reduction in the bacterial numbers when THP-1 cells were treated with EMB+NAC compared to EMB only. Data represent means ±SE from 6 trials. ***p<0.0005 when comparing infected macrophages treated with EMB+NAC to EMB lone treatment.

**Fig. 5C:**
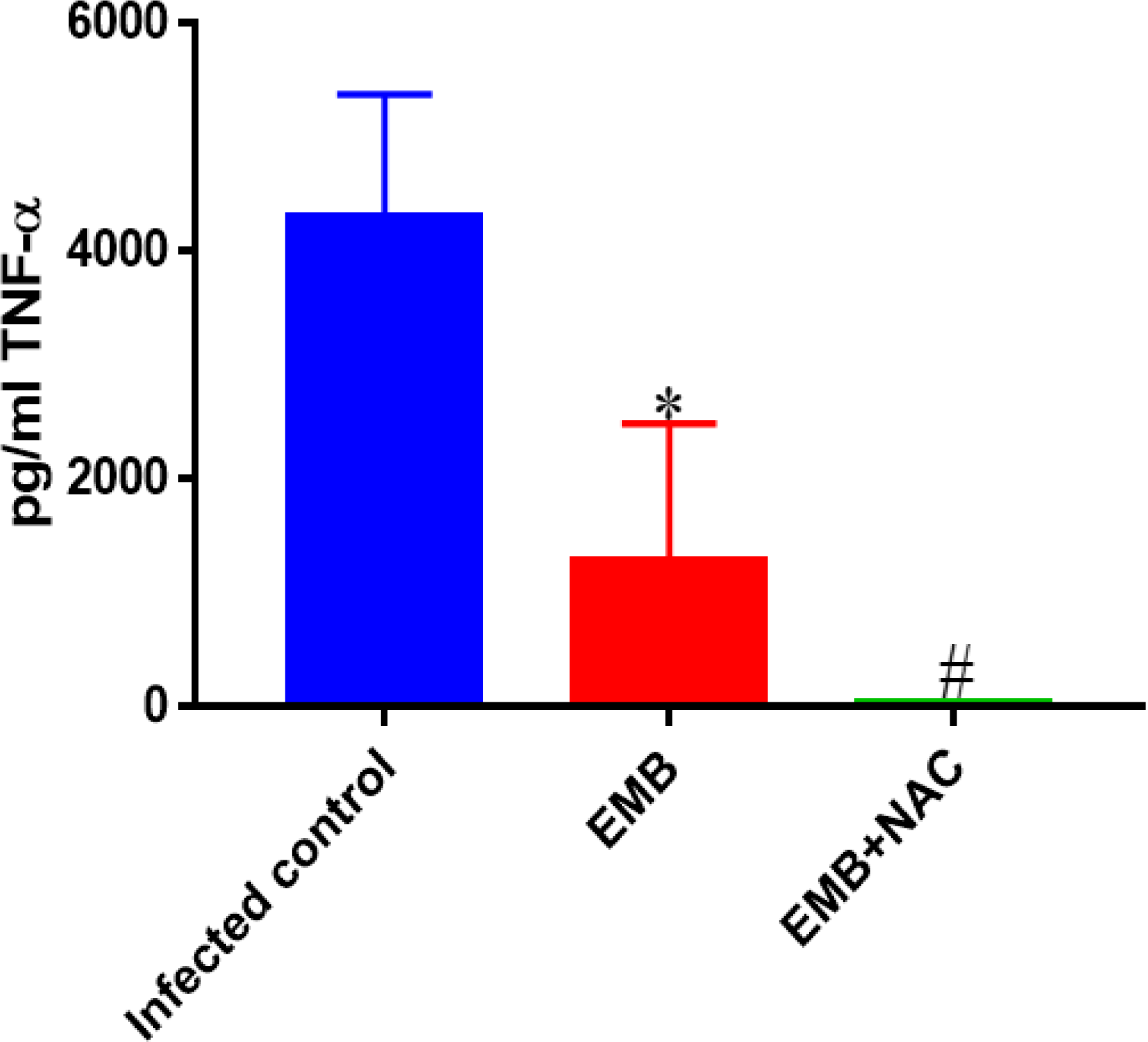
Assay of TNF-α in the supernatants from EMB and EMB + NAC-treated THP-1 cells. Assay of TNF-α was performed using an ELISA Ready-Set-Go kit from eBioscience. There was a significant decrease in the levels of TNF-**α** when samples were infected with *M*. *tb* and treated with EMB or EMB+NAC. Data represent means ±SE from 6 trials. *p<0.05 when comparing infected macrophages treated with EMB to infected sham controls. ^#^p<0.05 when comparing infected macrophages treated with EMB+NAC to treatment of infected sham controls.

**Fig. 5D:**
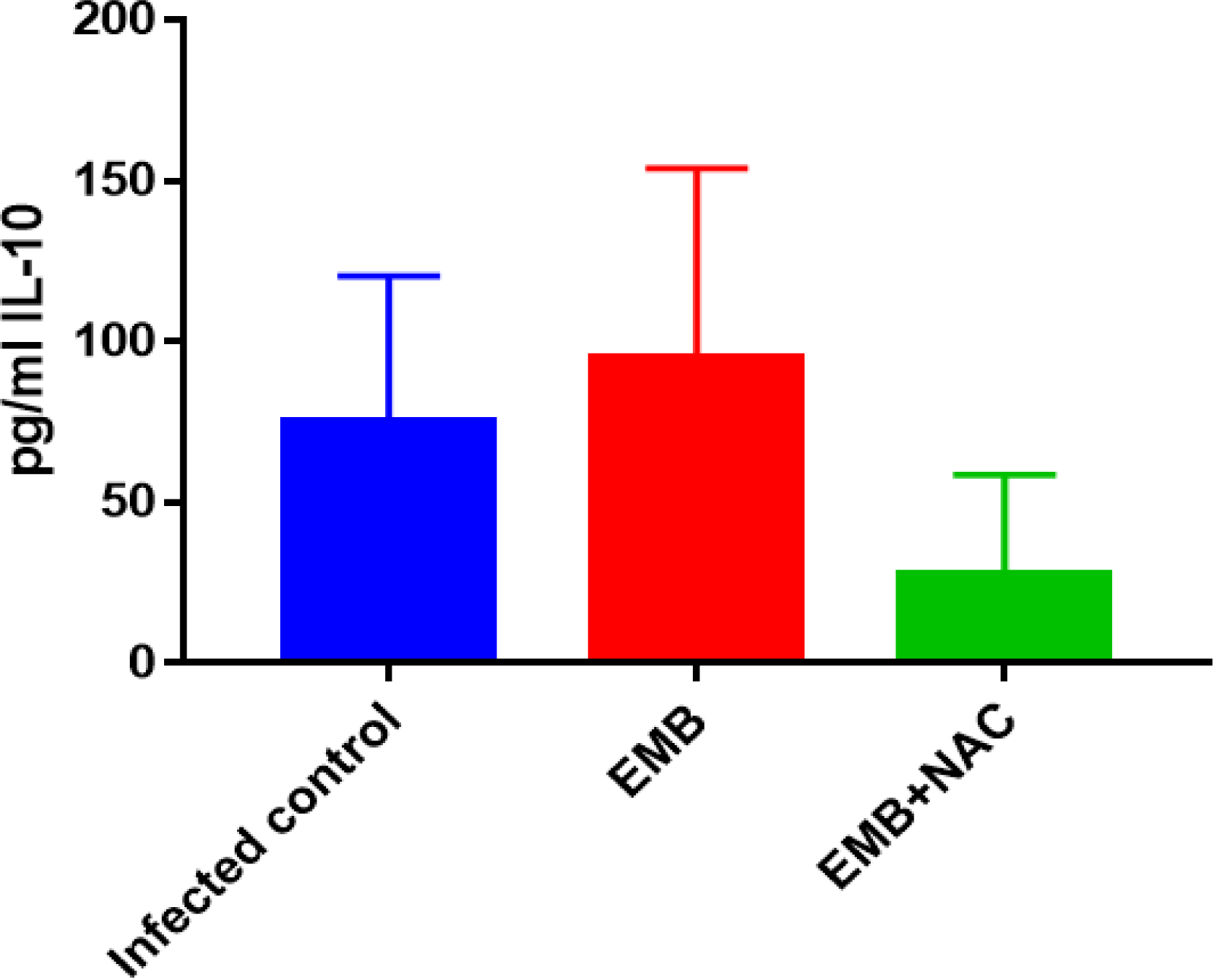
Assay of IL-10 in the supernatants from EMB and EMB + NAC-treated THP-1 cells. IL-10 levels were measured using an ELISA Ready-Set-Go kit from eBioscience. Although there was a decrease in levels of IL-10 when macrophages were infected with *M*. *tb* and treated with EMB+NAC, and a slight increase when samples were treated with EMB, these differences were not found to be statistically significant. Data represent means ±SE from 6 trials.

### Quantification of GSH levels, *M. tb* survival, TNF-α and IL-10 levels in M. *tb*-infected, *M*. *tb*-infected + PZA-treated and *M*. *tb*-infected with PZA+ NAC-treated THP-1 cells

PZA is an antibiotic that is normally given in combination with all four first-line antibiotics for the curative action against TB. To determine the effects of PZA and PZA+NAC in altering the intracellular survival of *M*. *tb* and production of GSH and cytokines, *M*. *tb* infected THP-1 cells were treated with stand-alone PZA (50 micrograms/ml) and the combination of PZA (50 micrograms/ml) and NAC (10mM) treatment. When compared to the infected sham control group the levels of GSH in PZA-treated macrophages were significantly elevated (Fig. 6A). There was further enhancement in the levels of GSH when *M*. *tb* infected macrophages were treated with PZA+ NAC (Fig. 6A). When compared to stand-alone PZA, the combination treatment of PZA+NAC led to a statistically significant decrease in CFUs of *M*. *tb* (Fig. 6B). PZA was more effective in lowering the bacterial load than both RIF (Fig. 4B) and EMB (Fig. 5B). Additionally, the treatment of PZA given in conjunction with NAC was as effective as the treatment of INH alone (Fig. 3B). TNF-α levels were quantified in the supernatants of *M. tb* infected macrophages in presence and absence of PZA sand PZA+NAC. The combined treatment of PZA+NAC exhibited a statistically significant decrease in levels of TNF-α compared to both the infected sham control and the stand-alone PZA categories (Fig. 6C). The levels of TNF-α after treatment with PZA was similar to that of EMB (Fig. 5C) and RIF (Fig. 4C). In comparison to sham control group, IL-10 levels were decreased in M. *tb* infected macrophages treated with PZA and PZA+NAC (Fig. 6D). Although not statistically significant, the PZA-treated macrophages showed a marked decrease in IL-10 levels compared to the sham control group. Importantly, treatment with PZA+NAC resulted in a statistically significant decrease in the levels IL-10, compared to the sham-control group (Fig. 6D). Furthermore, of all the first-line antibiotics tested, PZA treatment resulted in the lowest levels of IL-10 measured.

**Fig. 6A:**
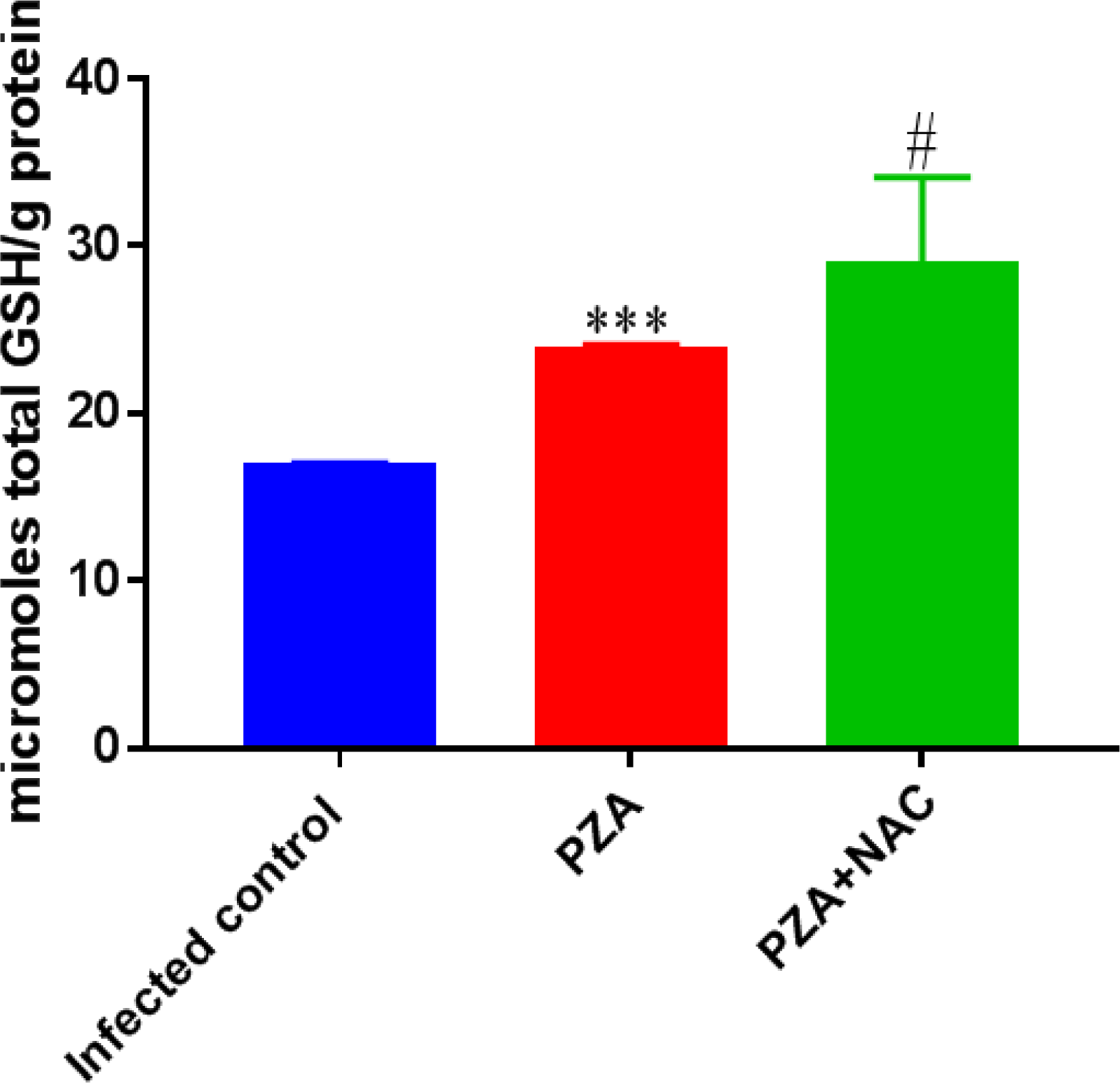
Levels of total GSH in *M*. *tb*-infected, *M. tb*-infected + PZA-treated and *M*. *tb*-infected + PZA + NAC-treated THP-1 cells. GSH assay was performed using a colorimetric assay kit from Arbor Assays. There was a significant increase in levels of GSH when infected macrophages were treated with PZA+NAC and PZA only. Data represent means ±SE from 6 trials. ^#^p<0.05 when comparing infected macrophages treated with PZA+NAC to infected sham controls. ***p<0.0005 when comparing infected macrophages treated with PZA to infected sham controls.

**Fig. 6B:**
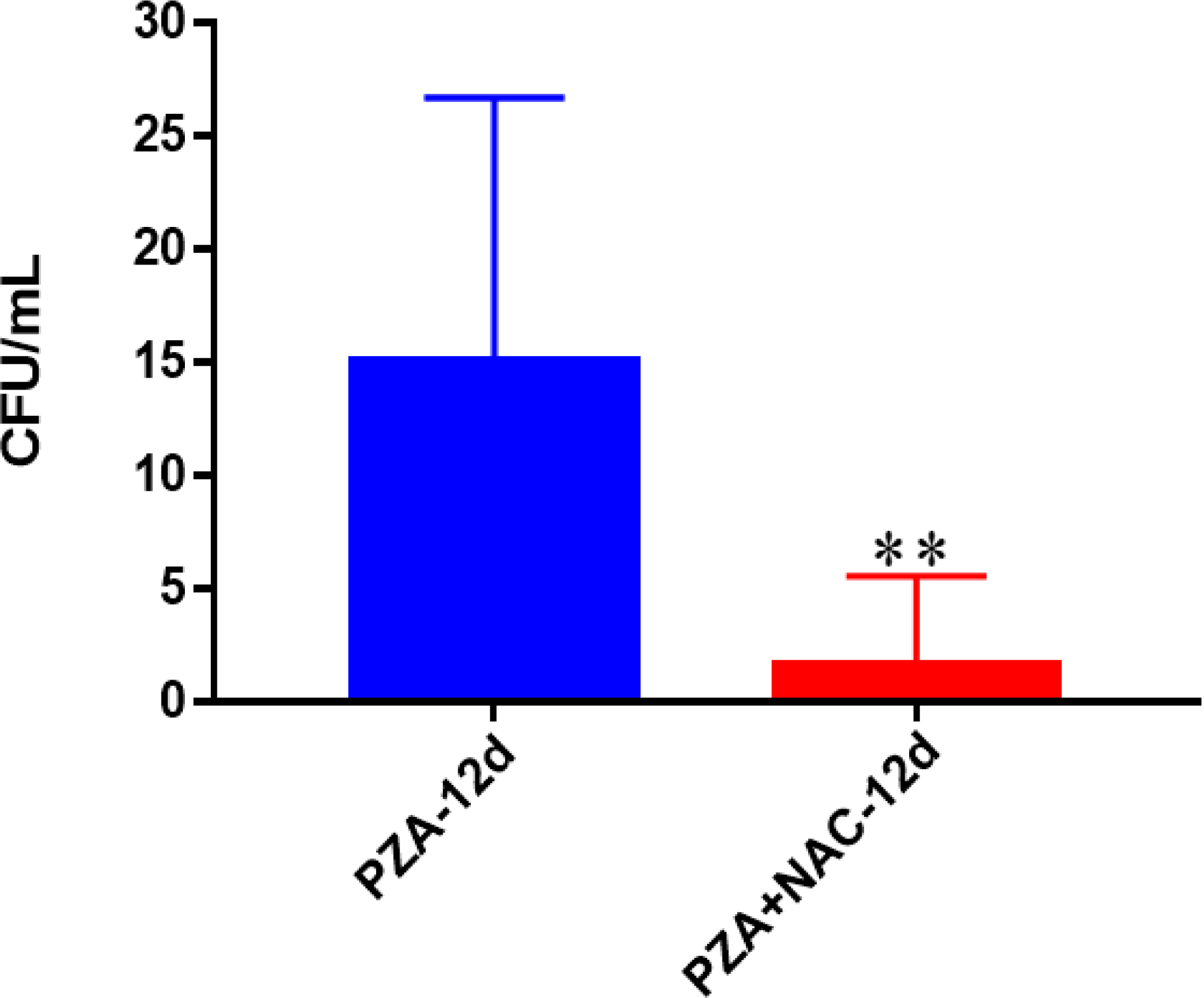
Survival of Erdman strain of *M*. *tb* inside PZA and PZA + NAC-treated THP-1 macrophages. THP-1 cells were cultured in a medium of RPMI and 10% FBS, and allowed to differentiate into macrophages by addition of PMA at a concentration of 10 ng/ml. There was a significant reduction of bacterial numbers when THP-1 cells were treated with PZA+NAC compared to PZA only. Data represent means ±SE from 6 trials. **p<0.005 when comparing infected macrophages treated with PZA+NAC to PZA only treatment.

**Fig. 6C:**
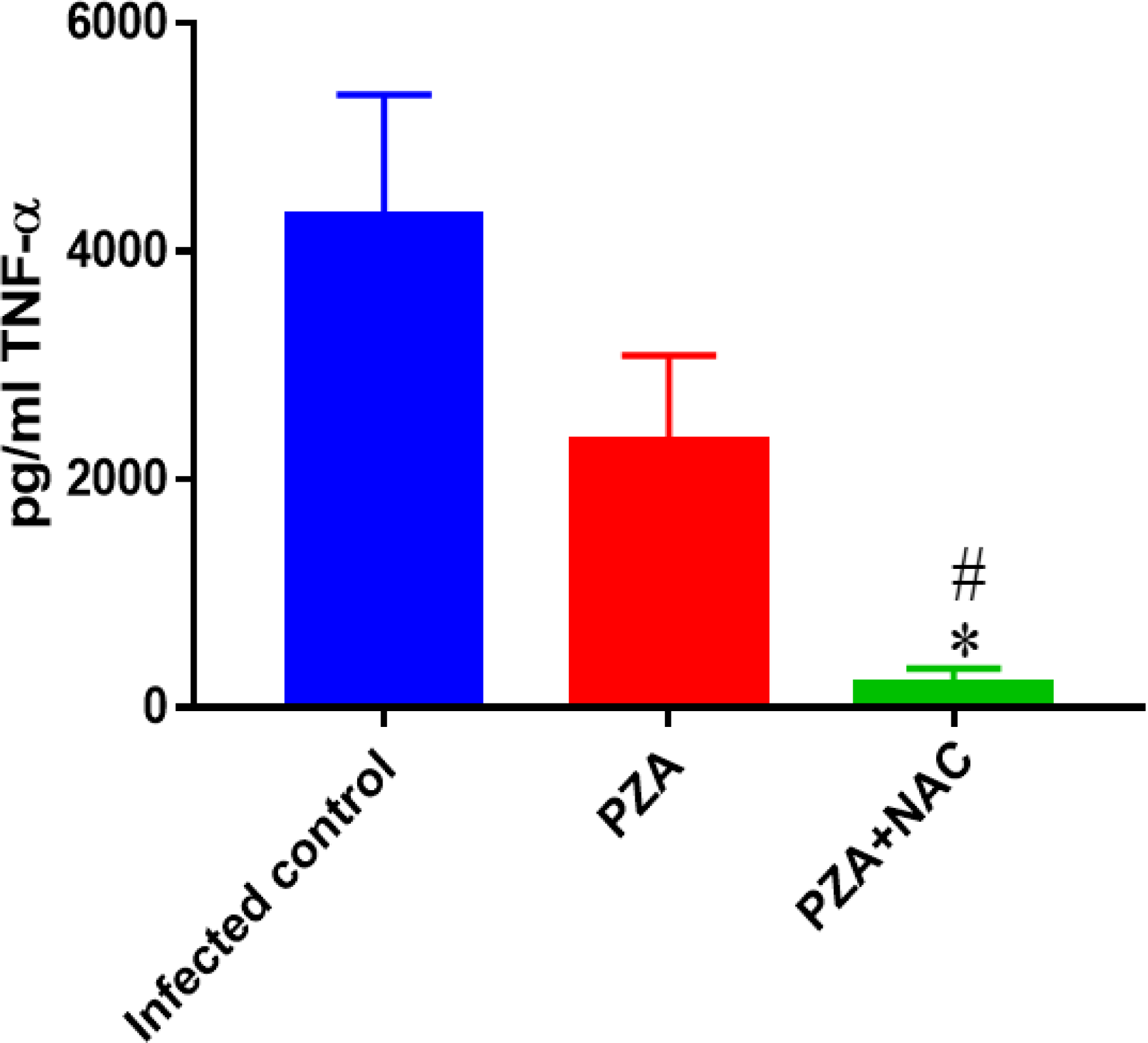
Assay of TNF-α in the supernatants from PZA and PZA + NAC-treated THP-1 cells. Assay of TNF-α was performed using an ELISA Ready-Set-Go kit from eBioscience. There was a significant decrease in the levels of TNF-α when macrophages were infected with *M*. *tb* and treated with PZA+NAC. Data represent means ±SE from 6 trials. *p<0.05 when comparing infected macrophages treated with PZA+NAC to infected sham controls. ^#^p<0.05 when comparing infected macrophages treated with PZA+NAC to treatment with PZA only.

**Fig. 6D:**
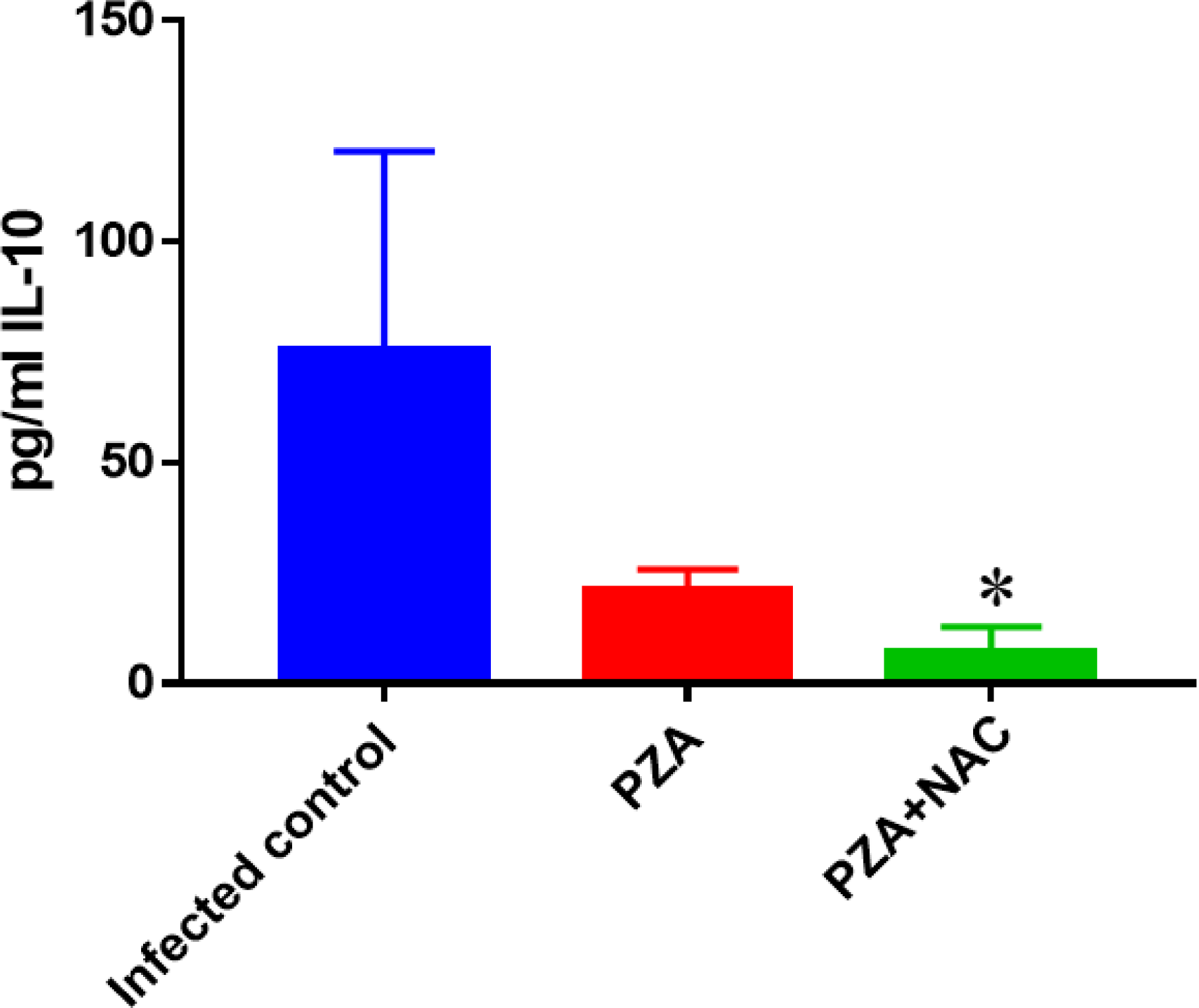
Assay of IL-10 in the supernatants from PZA and PZA + NAC-treated THP-1 cells. IL-10 assay was performed using an ELISA Ready-Set-Go kit from eBioscience. There was a significant decrease in the levels of IL-10 when macrophages were infected with *M. tb* and treated with PZA+NAC, and a slight decrease when samples were treated with PZA (not significant). Data represent means ±SE from 6 trials. *p<0.05 when comparing infected macrophages treated with PZA+NAC to infected sham control.

## Discussion

Immunocompromised individuals, such as HIV positive subjects, are increasingly susceptible to *M. tb* infection. Additionally, the rapid increase in the number of people living with drug resistant TB (DR-TB) and TB/HIV co-infection generates additional challenges for global targets of TB elimination. The standard six-month first-line antibiotic course of treatment consists of two phases: the intensive phase (the first two months) and the continuation phase (the last four months). However, treatment may differ for patients with extrapulmonary TB (TB outside of the lungs). During the intensive phase of standard treatment, patients take a daily combination of four medications: INH, RIF, PZA and EMB. These four drugs are referred to as first-line drugs and are off-patent and relatively inexpensive. However, the treatment of TB can become more complicated due to the development of drug-resistant and multidrug-resistant strains of *M. tb* [30]. Additionally, there are a variety of severe complications and side effects which can occur when taking these medications, especially when consuming the large dosage required to ensure proper elimination of a *M. tb* infection.

We previously reported that the virulent laboratory strain of *M. tb*, H37Rv is sensitive to physiological concentrations of GSH (5mM) when grown i*n vitro* (43, 44). We also found that enhancing the levels of GSH in human macrophages by treatment with NAC (10 or 20 mM) resulted in inhibition in the growth of intracellular H37Rv (43-46, 49). Thus, GSH has direct antimycobacterial activity, functioning as an effector molecule in innate defense against *M*. *tb* infection (43-49). These results unfold a novel and potentially important innate defense mechanism adopted by human macrophages to control *M*. *tb* infection (43-49). We also reported that GSH in combination with cytokines such as IL-2 and IL-12 enhances the functional activity of NK cells to inhibit the growth of M. *tb* inside human monocytes (47). We then demonstrated that GSH activates the functions of T lymphocytes to control *M. tb* infection inside human monocytes (48). These results indicate that GSH inhibits the growth of *M*. *tb* by both direct antimycobacterial effects as well as by enhancing the functions of immune cells (43-47). Finally, we demonstrated that the levels of GSH were significantly compromised in individuals with active pulmonary TB (49).

Therefore, we tested the synergistic effects of NAC (as a GSH precursor) and sub-optimal levels of individual first-line anti-TB drugs (INH, RIF, ETMB and PZA) in mediating control of *M. tb* infection inside THP-1 macrophages. The THP-1 cell line is an immortalized monocyte cell line, derived from the blood of a childhood case of acute monocytic leukemia [31, 32]. Each antibiotic was administered only once throughout the trial at its MIC. The MIC was administered to ensure that complete bacterial clearance was not achieved by the supplementation of the antibiotics alone.

Our results demonstrate that macrophages treated with the combination of suboptimal levels of each of the first-line antibiotics given in conjunction with NAC resulted in a significant reduction in the intracellular survival of *M. tb*, when compared to the administration of unaccompanied suboptimal levels of antibiotics (Fig. 3B, Fig. 4B, Fig. 5B, and Fig. 6B). In fact, complete clearance was observed for the INH category after the additional supplementation of NAC was added (Fig. 3B). These novel findings illustrate the synergistic effects of NAC/GSH and antibiotics in improving macrophages ability to control intracellular *M. tb* and suggests that GSH has suitable potential as an adjunct with the aforementioned first-line antibiotics in clearing a *M. tb* infection and aiding in the cessation of drug resistant strains of *M. tb*.

We observed a significant decrease in the levels of intracellular GSH in *M. tb*-infected macrophages compared to uninfected macrophages (Fig 1A). These results indicate that M. *tb* infection can cause intracellular GSH depletion, which in turn can promote *M*. *tb* survival and replication inside the host cells (Fig 1C). Furthermore, enhancing the levels of GSH in M. *tb*-infected macrophages by treatment with NAC (Fig 2A) resulted in significant reduction in the intracellular survival of *M. tb* (Fig 2B). Treatment of *M*. *tb*-infected macrophages with each of the first-line antibiotics resulted in restoration in the levels of GSH and a statistically significant (except of EMB*) increase when compared to the *M*. *tb*-infected sham control group (Fig. 3A, Fig. 4A, Fig. 5A, and Fig. 6A). These results indicate that use of antibiotics to limit *M*. *tb* burden in the macrophages enabled the host cells to restore the levels of GSH and improved their ability to combat the infection. Importantly, treatment of *M. tb*-infected macrophages with NAC in conjunction with each of the first-line antibiotics resulted in a statistically significant notable increase in the levels of GSH in all the categories (Fig. 3A, Fig. 4A, Fig. 5A, and Fig. 6A). The levels of GSH detected in macrophages treated with antibiotic in combination with NAC were consistently higher when compared sham control and macrophages treated with antibiotic alone (Fig. 3A, Fig. 4A, Fig. 5A, and Fig. 6A). Antibiotic treatment when given in conjunction with NAC resulted in decreased production of TNF-α and IL-10, by the macrophages. In the context of a *M. tb* infection, TNF-α is an inflammatory cytokine produced by macrophages and is particularly important in promoting the formation and maintenance of a granuloma, whereas IL-10 is an immunosuppressive cytokine which acts as negative regulator to the immune response associated with fighting of the infection [33-35]. TNF-α is a diverse cytokine, which has been shown to functionally aid in the formation and maintenance of a granuloma as well as play a critical role in host defense against *M. tb* in both acute phase and chronic phase of infection [36-38]. However, at high levels, TNF-α is also implicated as the source of many inflammatory and autoimmune diseases and has been shown to cause severe tissue damage when overexpressed in relation to a *M. tb* infection [39, 40]. When the combination of NAC and each of the first-line antibiotics was administered, the TNF-α levels for each category was significantly reduced from the extremely high levels seen among the infected sham control and presented closer to that of the uninfected group (Fig. 1C, Fig. 3C, Fig. 4C, Fig. 5C, and Fig. 6C). Our results indicate that NAC-treatment can modulate the levels of TNF-α in a manner that it is significant enough to maintain a healthy granuloma but cannot cause cellular damage. Increased levels of the cytokine IL-10 can dampen the effector responses against a *M. tb* infection by inhibiting phagosome-lysosome fusion within macrophages [40, 41, 42]. Although not statistically significant, a notable reduction in the levels of IL-10 was observed for every antibiotic category when administered with NAC (Fig. 3D, Fig. 4D, Fig. 5D, and Fig. 6D). Additionally, when NAC was supplemented unaided, a reduction of roughly double the magnitude of IL-10 was likewise detected (Fig. 2D). This data implies that NAC supplementation supports the immune responses in favor of eliminating a *M. tb* infection by reducing the levels of IL-10, allowing macrophage phagosome maturation and thus enhanced pathogen elimination. Our findings illustrate that in addition to the direct antimycobacterial effects, GSH-enhancement improved the ability of first-line antibiotics to limit intracellular *M*. *tb* infection and modulated the cytokine production by macrophages.

When all four antibiotics were administered together at their MICs, complete mycobacterial clearance was observed with and without the addition of NAC (data not shown). A similar trend was observed as with the supplementation of individual antibiotics, where a significant increase in levels of GSH was detected from the administration of all four antibiotics without NAC and a further increase once NAC was additionally added (data not shown). Likewise, a reduction in the levels of TNF-a, and IL-10 was observed among the administration of all four antibiotics without NAC supplementation and with statistical significance once NAC was also administered (data not shown).

Our findings highlight that GSH exhibits more physiological significance than just intracellular redox homeostasis, advocating that its enhancement aids in cytokine balance as well as augments the ability of first-line anti-TB drugs to clear a *M. tb* infection. Therefore, we believe that including GSH (NAC) in the antibiotic treatment of TB would not only limit cellular damage by means of redox balance, and subsequently reduce potential toxicity to the anti-TB medications but could possibly limit the required dosage necessary to cause complete bacterial clearance as well as help combat further emergences of DR-TB strains.

## Acknowledgments

The authors appreciate the funding support from Western University of Health Sciences and financial support of NSFC grants (No. 81272444 and 81472744) to conduct this study. Sincere thanks to Dr. Jozef Stec, Marshall B. Ketchum University for providing first-line antibiotics for this study.

## Author Contribution statement

Ruoqiong Cao – Maintained and Processed macrophages and mycobacteria for experiments, Preformed infection studies, Preformed ELISA assays, Preformed microscopy work.

Garrett Teskey – Wrote the abstract, methods, and discussion sections, Maintained and Processed macrophages and mycobacteria for experiments, Preformed infection studies, Preformed ELISA assays, Preformed microscopy work.

Hicret Islamoglu - Maintained and Processed macrophages and mycobacteria for experiments, Preformed infection studies, Preformed ELISA assays, Preformed microscopy work, contributed to writing the results section.

Rachel Abrahem - Preformed microscopy work, Wrote the results section.

Karo Gyurjian - Contributed to writing the abstract and introduction sections, Made media for bacterial quantification.

Li Zhong - Provided funding.

Vishwanath Venketaraman - Conceived the studies, Prepared figures, interpreted data, provided funding and prepared the manuscript.

## Additional information

The authors declare that there are **NO** competing interests for this publication.

## References

1. Smith I. Mycobacterium tuberculosis Pathogenesis and Molecular Determinants of Virulence. Clinical Microbiology Reviews. 2003;16(3):463–496.

2. World Health Organization (2017) Tuberculosis & Diabetes [Fact sheet]

3. Guirado E, Schlesinger LS, Kaplan G. Macrophages in Tuberculosis: Friend or Foe. Seminars in immunopathology. 2013;35(5):563–583.

4. Volkman HE, Clay H, Beery D, Chang JC, Sherman DR, Ramakrishnan L. Tuberculous granuloma formation is enhanced by a mycobacterium virulence determinant. PLOS Biol. 2004 Nov;2(11): e367.

5. Guirado E, Mbawuike U, Keiser TL, Arcos J, Azad AK, Wang SH, Schlesinger LS: Characterization of host and microbial determinants in individuals with latent tuberculosis infection using a human granuloma model. mBio 2015, 6(1): e02537–02514.

6. Domingo-Gonzalez R, Prince O, Cooper A, Khader SA. Cytokines and Chemokines in *Mycobacterium tuberculosis* Infection. Microbiol Spectr. 2016 Oct;4(5).

7. Ehlers S, Schaible UE. The Granuloma in Tuberculosis: Dynamics of a Host–Pathogen Collusion. Frontiers in Immunology. 2012;3:411.

8. Dorman SE, Chaisson RE: From magic bullets back to the magic mountain: the rise of extensively drug-resistant tuberculosis. Nature medicine 2007, 13(3):295–298.

9. Ormerod LP: Multidrug-resistant tuberculosis (MDR-TB): epidemiology, prevention and treatment. British Medical Bulletin 2005, 73-74(1):17–24.

10. Udwadia ZF. MDR, XDR, TDR tuberculosis: ominous progression. Thorax. 2012 Apr; 67(4):286–8.

11. Horne DJ, Spitters C, Narita M. Experience with Rifabutin Replacing Rifampin in the Treatment of Tuberculosis. The International Journal of Tuberculosis and Lung Disease. 2011;15(11):1485–i.

12. Leibert E, Rom WN. New drugs and regimens for treatment of TB. Expert review of anti-infective therapy. 2010;8(7):801–813.

13. Shenoi S, Heysell S, Moll A, Friedland G. Multidrug-resistant and extensively drug-resistant tuberculosis: consequences for the global HIV community. Current opinion in infectious diseases. 2009;22(1):11–17.

14. Arbex MA, Varella Mde C, Siqueira HR, Mello FA. Antituberculosis drugs: drug interactions, adverse effects, and use in special situations. Part 1: first-line drugs. J Bras Pneumol. 2010 Sep-Oct;36(5):626-40. Review.

15. Thwaites G, Auckland C, Barlow G, Cunningham R, Davies G, Edgeworth J, Greig J, Hopkins S, Jeyaratnam D, Jenkins N et al: Adjunctive rifampicin to reduce early mortality from Staphylococcus aureus bacteraemia (ARREST): study protocol for a randomised controlled trial. Trials 2012, 13(1):241.

16. Ramappa V, Aithal GP. Hepatotoxicity Related to Anti-tuberculosis Drugs: Mechanisms and Management. Journal of Clinical and Experimental Hepatology. 2013;3(1):37–49.

17. FDA News Release; United States Food and Drug Administration: Silver Spring, MD, dec 31, 2012.

18. Brigden G, Hewison C, Varaine F. New developments in the treatment of drug-resistant tuberculosis: clinical utility of bedaquiline and delamanid. Infection and Drug Resistance. 2015;8:367–378.

19. Bavarsad Shahripour R, Harrigan MR, Alexandrov AV. N-acetylcysteine (NAC) in neurological disorders: mechanisms of action and therapeutic opportunities. Brain and Behavior. 2014;4(2):108–122.

20. Forman HJ, Zhang H, Rinna A. Glutathione: Overview of its protective roles, measurement, and biosynthesis. Molecular aspects of medicine. 2009;30(1-2):1–12.

21. Saing T, Lagman M, Castrillon J, Gutierrez E, Guilford FT, Venketaraman V. Analysis of glutathione levels in the brain tissue samples from HIV-1-positive individuals and subject with Alzheimer’s disease and its implication in the pathophysiology of the disease process. BBA Clin. 2016 May 29;6:38–44.

22. Allen M, Bailey C, Cahatol I, Dodge L, Yim J, Kassissa C, Luong J, Kasko S, Pandya S, Venketaraman V. Mechanisms of Control of Mycobacterium tuberculosis by NK Cells: Role of Glutathione. Front Immunol. 2015 Oct 5;6:508.

23. Ly J, Lagman M, Saing T, Singh MK, Tudela EV, Morris D, Anderson J, Daliva J, Ochoa C, Patel N, Pearce D, Venketaraman V. Liposomal Glutathione Supplementation Restores TH1 Cytokine Response to Mycobacterium tuberculosis Infection in HIV-Infected Individuals. J Interferon Cytokine Res. 2015 Nov;35(11):875–87.

24. Morris D, Ly J, Chi PT, Daliva J, Nguyen T, Soofer C, Chen YC, Lagman M, Venketaraman V. Glutathione synthesis is compromised in erythrocytes from individuals with HIV. Front Pharmacol. 2014 Apr 11; 5:73.

25. Morris D, Nguyen T, Kim J, Kassissa C, Khurasany M, Luong J, Kasko S, Pandya S, Chu M, Chi PT, Ly J, Lagman M, Venketaraman V. An elucidation of neutrophil functions against Mycobacterium tuberculosis infection. Clin Dev Immunol. 2013;2013:959650.

26. Morris D, Gonzalez B, Khurasany M, Kassissa C, Luong J, Kasko S, Pandya S, Chu M, Chi PT, Bui S, Guerra C, Chan J, Venketaraman V. Characterization of dendritic cell and regulatory T cell functions against Mycobacterium tuberculosis infection. Biomed Res Int. 2013;2013:402827.

27. Lagman M, Ly J, Saing T, Kaur Singh M, Vera Tudela E, Morris D, Chi PT, Ochoa C, Sathananthan A, Venketaraman V: Investigating the causes for decreased levels of glutathione in individuals with type II diabetes. PLOS one 2015, 10(3):e0118436.

28. Amaral EP, Conceicao EL, Costa DL, Rocha MS, Marinho JM, Cordeiro-Santos M, D’Imperio-Lima MR, Barbosa T, Sher A, Andrade BB: N-acetyl-cysteine exhibits potent anti-mycobacterial activity in addition to its known anti-oxidative functions. BMC microbiology 2016, 16(1):251.

29. Daoud AK, Tayyar MA, Fouda IM, Harfeil NA. Effects of diabetes mellitus vs. in vitro hyperglycemia on select immune cell functions. J Immunotoxicol. 2009 Mar;6(1):36–41.

30. Palomino JC, Martin A. Drug Resistance Mechanisms in *Mycobacterium tuberculosis*. Antibiotics. 2014;3(3):317–340.

31. Bosshart H, Heinzelmann M. THP-1 cells as a model for human monocytes. Annals of Translational Medicine. 2016;4(21):438.

32. Tsuchiya S, Yamabe M, Yamaguchi Y, et al. Establishment and characterization of a human acute monocytic leukemia cell line (THP-1). Int J Cancer 1980;26:171-6. 10.

33. Shrivastava P, Bagchi T. IL-10 modulates in vitro multinucleate giant cell formation in human tuberculosis. PLOS One. 2013 Oct 17;8(10):e77680.

34. Tanaka T, Narazaki M, Kishimoto T. IL-6 in Inflammation, Immunity, and Disease. Cold Spring Harbor Perspectives in Biology. 2014;6(10):a016295. doi:10.1101/cshperspect.a016295.

35. Mootoo A, Stylianou E, Arias MA, Reljic R. TNF-alpha in tuberculosis: a cytokine with a split personality. Inflamm Allergy Drug Targets. 2009 Mar;8(1):53–62.

36. Chan J, Flynn J. The immunological aspects of latency in tuberculosis. Clin Immunol. 2004 Jan;110(1):2-12. Review.

37. Dorhoi A, Kaufmann SH. Tumor necrosis factor alpha in mycobacterial infection. Semin Immunol. 2014 Jun;26(3):203-9. doi:10.1016/j.smim.2014.04.003. Epub 2014 May 10. Review.

38. Flynn JL, Chan J. What’s good for the host is good for the bug. Trends Microbiol. 2005 Mar;13(3):98-102. Review.

39. Dreher D, Nicod LP (2002) Dendritic cells in the mycobacterial granuloma are involved in acquired immunity. Am J Respir Crit Care Med 165: 1577–1578.

40. Esposito E, Cuzzocrea S. TNF-alpha as a therapeutic target in inflammatory diseases, ischemia-reperfusion injury and trauma. Curr Med Chem. 2009;16(24):3152–67.

41. Wolff SP, Dean RT: Glucose autoxidation and protein modification. The potential role of ‘autoxidative glycosylation’ in diabetes. The Biochemical journal 1987, 245(1):243–250.

42. O’Leary S, O’sullivan MP, Keane J. IL-10 blocks phagosome maturation in mycobacterium tuberculosis-infected human macrophages. Am J Respir Cell Mol Biol. 2011 Jul;45(1):172–80.

43. Venketaraman V, Dayaram YK, Talaue MT, Connell ND. 2005. Glutathione and nitrosoglutathione in macrophage defense against Mycobacterium tuberculosis. Infect Immun. 73(3):1886–9.

44. Dayaram, Y. K., Talaue, M. T., Connell, N. D., and Venketaraman, V. 2006. Characterization of a glutathione metabolic mutant of Mycobacterium tuberculosis and its resistance to glutathione and nitrosoglutathione. J Bacteriology. 188 (4):1364–72.

45. Morris D, Guerra C, Khurasany M, Guilford F, Saviola B, Huang Y and Venketaraman V. 2013. Glutathione supplementation improves macrophage functions in HIV. Journal of Interferon and cytokine research. 33 (5): 270–9.

46. Morris D, Guerra C, Donohue C, Oh H, Khurasany M, Venketaraman V. 2012. Unveiling the mechanisms for decreased glutathione in individuals with HIV infection. Clin Dev Immunol. 2012:734125.

47. Guerra C, Johal K, Morris D, Moreno S, Alvarado O, Gray D, Tanzil M, Pearce D, Venketaraman V. 2012. Control of Mycobacterium tuberculosis growth by activated natural killer cells. Clin Exp Immunol. 168(1):142–52.

48. Guerra G, Morris D, Gray D, Tanzil M, Sipin A, Kung S, Guilford F, Khasawneh F and Venketaraman V. 2011. Adaptive immune responses against Mycobacterium tuberculosis infection in healthy and HIV infected individuals. PLoS One. 2011. 6(12): e28378.

49. Venketaraman, V; Millman, AC; Salman, M; Swaminathan, S; Goetz, M; Lardizabal, A; Hom, D; and Connell, N. D. 2008. Glutathione levels and immune responses in tuberculosis patients. Microbial pathogenesis. 44: 255–261.

